# Recent hybridization and allopolyploidy reprogrammed *Spartina* microRNA expression under xenobiotic induced stress

**DOI:** 10.1101/2019.12.13.875138

**Authors:** Armand Cavé-Radet, Armel Salmon, Loup Tran Van Canh, Richard L. Moyle, Lara-Simone Pretorius, O. Lima, Malika L. Ainouche, Abdelhak El Amrani

## Abstract

Xenobiotic detoxification is a common trait of all living organisms, necessary for developmental plasticity and stress tolerance. The gene set involved in this biological process is dubbed the xenome (*i.e.* involved in drug metabolism in mammals, degradation of allelochemicals and environmental pollutants by bacteria and plant communities). Recently, we found that allopolyploidy increased tolerance to xenobiotics (phenanthrene) in *Spartina*. To decipher the molecular mechanisms underlying this process, we examined how interspecific hybridization and genome doubling impact miRNAs expression under xenobiotic induced stress. In this work we used a deep sequencing approach, and analyzed the parental species *S. alterniflora* and *S. maritima*, their F1 hybrid *S. x townsendii* and the allopolyploid *S. anglica* under phenanthrene exposure. We found that hybridization and genome doubling reprogrammed a myriad of miRNAs under phenanthrene-induced stress. Hence, to identify the master miRNAs involved in phenanthrene tolerance, we performed experimental functional validation of phenanthrene-responsive Spar-miRNAs using Arabidopsis T-DNA mutant lines inserted in homologous MIR genes, 39 knock out T-DNA *Arabidopsis* mutants, tagged in the most conserved miRNAs genes in vascular plants were screened. Development of MIR159 and MIR156 mutants was significantly affected under phenanthrene-induced stress. Subsequently, we performed *in planta* experimental validation to confirm the interaction between these miRNAs and their targets. These analyses suggest that MIR159 and MIR156 regulatory modules were targeted to induce the xenome relaxation and impact developmental plasticity responses in phylogenetically distant species under xenobiotic-induced stress.

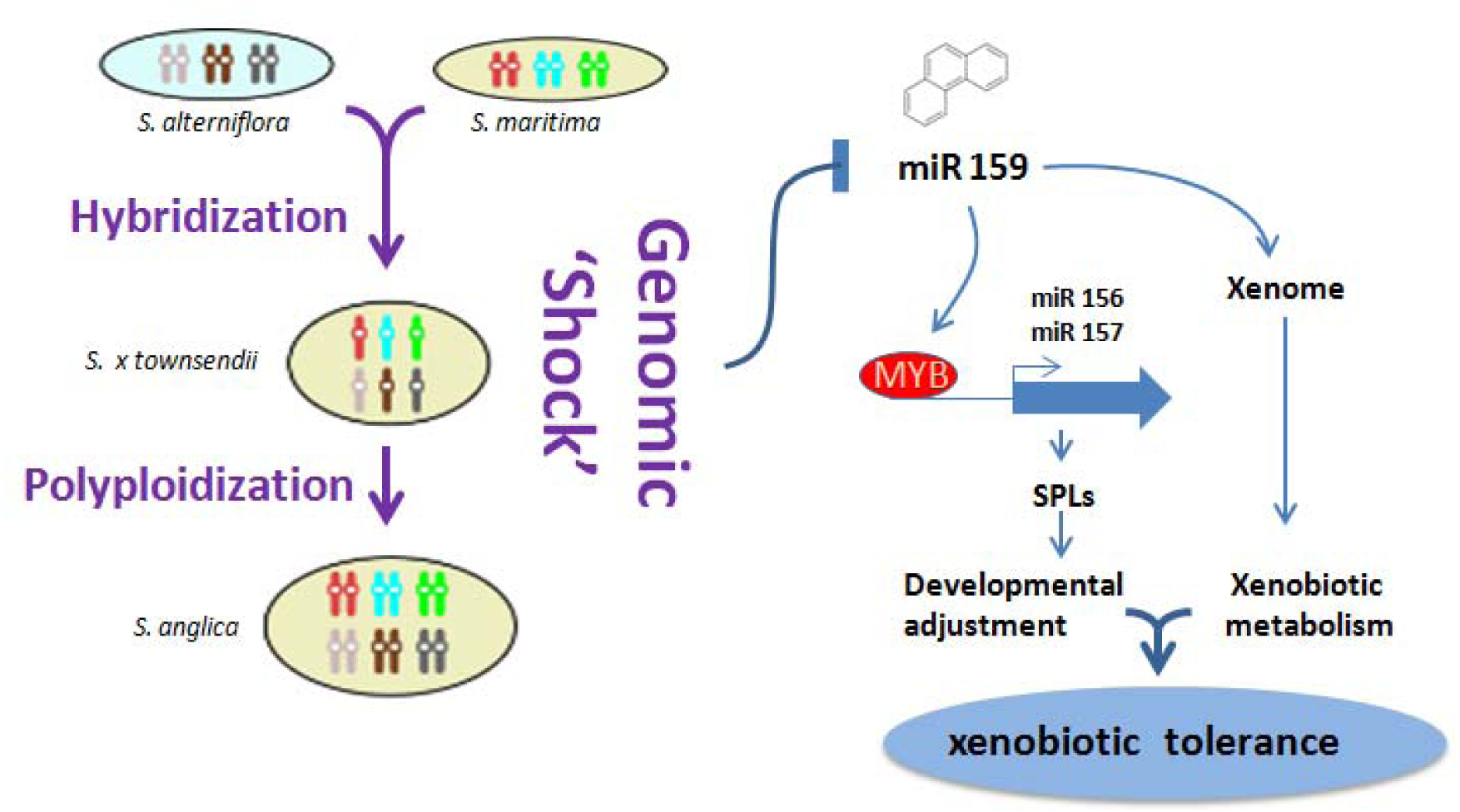
Graphical abstract

## INTRODUCTION

Xenobiotic detoxification is an adaptive trait encountered in all living organisms. This process is a determinant feature used in a myriad of environmental situations, such as the selection of ecological partners mediated by allelochemicals (Dong *et al*., 2016), drug metabolism in mammals (Yang *et al*., 2018), fungal degradation of various recalcitrant organic matter and more generally potential toxic compounds resulting from wood degradation (Morel *et al*., 2013), and finally degradation and dissipation of pollutants by bacteria and plant communities (El Amrani *et al*., 2015). However, knowledge of xenobiotic detoxification pathways in higher plants is still fragmentary. Among the organic xenobiotic compounds, PAHs (Polycyclic Aromatic Hydrocarbons) represent widespread pollutants, alarming by their environmental toxicity with ecological and public health issues. Physiological impact of PAHs on plant physiology is well documented, mainly in the plant model *Arabidopsis* (Alkio *et al*., 2005; Liu *et al*., 2009; Shiri *et al*., 2015; Dumas *et al*., 2016). Absorption of xenobiotics by plant roots occurs through passive or active transport mechanisms (Pilon-Smits, 2005; Zhan *et al*., 2012). Once in plant tissues, xenobiotics may be translocated from root to shoot tissues by xylem vascular elements where they are accumulated, degraded, and/or compartmented (Gao & Zhu, 2004; Pilon-Smits, 2005). In plants, PAHs reduce growth and development by disrupting major metabolic functions like photosynthesis. Depending on plant tolerance abilities, PAHs may induce necrosis and promote oxidative stress by ROS (Reactive Oxygen Species) production, that may limit plant tolerance (Alkio *et al*., 2005; Shiri *et al*., 2015; Dumas *et al*., 2016).

The concept of the xenome emerged to characterize all genes involved in xenobiotic detoxification (Edwards *et al*., 2005, 2011; Taguchi *et al*., 2010; El Amrani *et al*., 2015; Dumas *et al*., 2016). It involves transport, signaling and sensing genes, and several gene families such as α/β hydrolases, cytochromes P450 (*CYPs*), glycosyltransferases (*GTs*) and malonyltransferases involved in xenobiotic conjugation and/or biotransformation. In addition, ATP-binding cassette transporters (ABC) may be responsible for xenobiotic conjugates transport for vacuolar storage, and several phytohormones like ethylene or jasmonic acid seem to be involved in PAH responses (Weisman *et al*., 2010). Transcriptomic analyses (using CATMA and ATH1 microarray profiling) in *Arabidopsis* under phenanthrene-induced stress (used as a model PAH) revealed significant up and down-regulations of xenome expression patterns in previously cited gene families (Weisman *et al*., 2010; Dumas *et al*., 2016), which highlighted candidate genes involved in primary responses to xenobiotics (*e.g*. signaling, sensing) and molecular mechanisms acting on putative xenobiotic metabolization. Recently, Alvarez *et al*. (2018) expanded the xenome to non-model plants from the *Spartina* lineage (Poaceae). In their study, they performed transcriptomic analyses on *Spartina alterniflora* populations which experienced crude oil exposure following the *Deepwater Horizon* oil spill. Comparisons between stress-responsive genes reported in *Arabidopsis* under phenanthrene *in vitro* experiments were performed, but little overlap between candidate xenome genes was detected, suggesting that molecular responses to xenobiotics may differ between phylogenetically distant species.

Most *Spartina* species are salt marsh plants, colonizing estuaries, coastal mudflats, or intertidal areas. *Spartina* species are characterized by recurrent hybridization and genome doubling events (Ainouche *et al*., 2009, 2012). In *Spartina*, a recent allopolyploidization event offers a unique opportunity to study the respective impacts of hybridization and genome doubling on structural and functional genome dynamics (Salmon *et al*., 2005; Parisod *et al*., 2009; Chelaifa *et al*., 2010b; Ferreira de Carvalho *et al*., 2017). *Spartina maritima* (paternal parent) and *Spartina alterniflora* (maternal parent) hybridized during the end of the 19^th^ century in islands of southern England and led the formation of the F1 hybrid *Spartina x townsendii*. Whole genome doubling of *S. x townsendii*, probably by the union of unreduced gametes resulted in the new allopolyploid species *Spartina anglica* (Hubbard, 1968). In *Spartina*, high tolerance abilities to stress such as salinity, anoxia and heavy-metals are reported (Anderson & Treshow, 1980; Thompson, 1991; Castillo *et al*., 2000; Hacker *et al*., 2001; Wang *et al*., 2006).

In allopolyploids, the genomic ‘shock’ resulting from genome merger and redundancy is known to induce important changes on structural and functional genome dynamics (Wendel, 2015; Wendel *et al*., 2018). Such changes may involve new gene expression patterns in the hybrids and/or the allopolyploids compared to their parents, and result in new phenotypes that may impact the adaptive potential of the newly formed species (Doyle *et al*., 2008). In this perspective, abilities to colonize vacant niches or stressful environments are frequently reported in polyploids (Wood *et al*., 2009). In their habitats, *Spartina sp.* are exposed to environmental pollution from anthropogenic activities like oil spills (Lin & Mendelssohn, 2012; Lin *et al*., 2016; Robertson *et al*., 2017), which include organic pollutants compounds from the PAH family. Interestingly, resilience abilities reported in *S. alterniflora* populations exposed to crude oil suggest efficient tolerance abilities to such environmental pollutants (Wu *et al*., 2012; Alvarez *et al*., 2018), that could be used for green remediation. Recently, comparative physiological analyses of phenanthrene (one of major PAHs in crude oil; Liu *et al*., 2012) treated *Spartina* leaves between parental species and *S. anglica* reported enhanced tolerance abilities to PAHs following allopolyploidy (Cavé-Radet *et al*., 2019b).

Owing to their central function, miRNAs are actively explored in various plant lineages, which most likely contributes to explain how gene networks are regulated. With the development of miRNA microarrays and the broad access to NGS technologies, studies of miRNA regulation mechanisms expanded in plant sciences. Various studies explored miRNA functions in molecular mechanisms of stress tolerance (Khraiwesh *et al*., 2012; Sunkar *et al*., 2012; Zhang, 2015; Shriram *et al*., 2016) highlighting their major regulatory role. For example, their impact in response to environmental abiotic stress like drought or salinity affecting plant growth, development and productivity were studied in crop species of agronomic interest such as maize (Li *et al*., 2013; Fu *et al*., 2017), wheat (Feng *et al*., 2017; Bakhshi *et al*., 2017) or tomato (Liu *et al*., 2017). Down or up-regulations of miRNAs under stress respectively up and down-regulate target genes expression, impacting the functionality of positive and negative stress tolerance regulators.

In order to decipher the molecular mechanisms underlying xenobiotic tolerance abilities, and to unravel the effects of allopolyploidization under such stress, we focused on miRNAs and their impact in the regulation of stress-responsive genes. Here, we hypothesized that miRNAs expression in response to stress following hybridization or genome doubling events in *Spartina* may be crucial in species tolerance abilities. We performed small RNA-Seq and RNA-Seq in *Spartina* species exposed to phenanthrene for 5 days, using leaf tissues sampled in *S. alterniflora*, *S. maritima*, *S. x townsendii* and *S. anglica*. Based on *Spartina* miRNAs (Spar-miRNAs) annotations (Cavé-Radet *et al*., 2019a), we conducted differential expression analyzes, in order to identify candidate phenanthrene-responsive miRNAs and mRNAs. We performed experimental functional validation of conserved phenanthrene-responsive Spar-miRNAs using *Arabidopsis* T-DNA mutant lines inserted in homologous *MIR* genes. In addition, *in planta* interactions between homologous stress-responsive between homologous phenanthrene-responsive miRNAs and their target genes in *Arabidopsis* were validated using dual luciferase assay strategy (Moyle *et al*., 2017). The reported results bring new insights in understanding the role of miRNA expression in response to PAHs, and the consequences of allopolyploidization in xenobiotic-stress tolerance.

## MATERIAL AND METHODS

### 1. Plant material, phenanthrene treatments and sequencing

We collected plants between 2012 and 2015 in their natural habitats along coastlines of France and England. The parental species were sampled at Le Faou (Finistère, France) for *S. alterniflora*, and in Brillac-Sarzeau and Le Hezo (Morbihan, France) for *S. maritima*. We collected the homoploid hybrid *S. x townsendii* in Hythe (Kent, England), and the allopolyploid *S. anglica* was sampled at La Guimorais (Ille-et-Vilaine, France). Plants were maintained the greenhouse under controlled environment until experiments.

Salt marsh sediment from the sampling site of *S. anglica* was rinsed in order to reduce the salt content. Once dried, we sieved substrate through a 5mm grid. Then, we relied on the experimental design described by Hong *et al*. (2015) and spiked 150 g of sediment with 1.950 g of phenanthrene (99%, Sigma-Aldrich) diluted in absolute ethanol (100 mM). We performed controls following the same procedure by adding identical volume (109.4 mL) of absolute ethanol without the xenobiotic. After total evaporation of ethanol for one day, we mixed vigorously these sediments by hand with an additional volume of air-dried sediment (12.85 kg) to reach an average final concentration of 150 mg phenanthrene.kg^−1^ in the polluted substrate. We supplemented both polluted and control sediments with sterile vermiculite (volume 1:3) for a better substrate aeration. Then, collected *Spartina* plants were potted (1L) with treated and control substrates. We performed three biological replicates for each species and condition (phenanthrene-treated and control), and placed plants in a phytotronic chamber with a light/dark regime of 16/8h with an average ambient temperature of 20°C, watered each day with 150 mL of half-strength Hoagland’s nutrient solution (Arnon & Hoagland, 1950).

After 5 days, leaves were harvested, immediately frozen in liquid nitrogen, and kept at -80°C. We performed RNA extraction with TRIzol® reagent (Sigma-Aldrich) according to the method adapted by Chelaifa *et al*. (2010a,b). In total, we extracted 24 samples corresponding to three biological replicates per species and treatment for sequencing. Small RNA-Seq and data preprocessing were performed as previously described in Cavé-Radet *et al*. (2019a), on 16 libraries (two of the three biological replicates performed). Sequencing was performed in single end, on a NextSeq500 (Illumina®) sequencer (1×50 pb, 20 million reads). RNA-Seq was performed for all 24 samples. Libraries were prepared using 1 µg of total RNA, with the « NEBNext^®^ Ultra™ II DNA Library Prep Kit » kit (Illumina®). Paired-end sequencing was performed in on a HiSeq1500 (Illumina®) sequencer (2×100 pb, 20 million de reads, GEH platform, Biogenouest, UR1).

### 2. *In silico* differential expression analysis

The annotation of miRNAs and their differential expression were processed as described in Cavé-Radet *et al*. (2019a). Reads were normalized with EDAseq (Risso *et al*., 2011) and differential expression tested with DESeq2 comparisons (Love *et al*., 2014). We processed both intraspecific comparisons (control *vs.* phenanthrene-treated) and interspecific pairwise comparisons (*i.e.* between *S. townsendii, S. anglica*, the parental species, and the Mid-Parent Value: MPV, representing an *in silico* average of parental expression) under phenanthrene. Impacts of hybridization and genome doubling were identified comparing expression patterns between the MPV against *S. x townsendii*, and between the F1 hybrid against *S. anglica*, respectively. Significantly DE miRNAs were identified according to Bonferroni adjusted p.values < 0.05.

From RNA-Seq data, we analyzed the expression of 9,714 contigs selected among putative miRNA target genes in *Spartina* species from Cavé-Radet *et al*. (2019a). *Spartina* transcripts were assembled under phenanthrene induced stress following the pipeline used for *Spartina* reference transcriptome assemblies (Boutte *et al*., 2016). Sequenced reads were mapped to *Spartina* contigs using Bowtie2 (Langmead & Salzberg, 2012) with the following parameters: --local -N 1 --score-min G,52,8 --no-contain --no-overlap --no-discordant. The number of mapped reads to each contig was extracted using Samtools idxstats command (Li *et al*., 2009) for each sequencing library, and combined into a counting matrix. Differential expression analyses were performed using the same pipeline (EDAseq and DESeq2) and the same comparisons as described for miRNAs.

### 3. Quantitative analysis of miRNAs by RT-qPCR

RT-qPCR of miRNAs in *Spartina* were performed using the poly(T) adaptor method (Shi *et al*., 2012), from the same RNA extracts used for deep sequencing. After DNase treatments of RNA extracts using the Turbo DNA-free kit (Ambion, Life technologies), cDNAs were produced by poly(A)-tailing (Poly(A) tailing kit, Ambion) and reverse transcription (SuperScript™ III Reverse Transcriptase kit, Invitrogen, Life technologies) using a universal poly(T) adaptor primer. PCR amplifications were monitored with a reverse primer annealing with the universal poly(T) adaptor sequence of cDNA. Forward primers used for specific miRNA amplifications were designed manually, according to mature miRNA sequences. We checked primer specificity to avoid any over-amplification related to the high similarity between miRNA sequences from the same family, and selected primers with Tm around 55°C estimated by OligoAnalyzer 3.1 (https://eu.idtdna.com/calc/analyzer). The designed primers are listed in Supplemental Table S1.

In total, we obtained 24 cDNA samples corresponding to three biological replicates per species and treatment for RT-qPCR. We normalized cDNA libraries at 3 ng.µL^−1^, and performed reactions in a final volume 20 µL. Reactions were carried out with 10 µL of SYBR Green Master Mix (Thermo Fisher), 5 ng of cDNA, forward and universal reverse primers (0.8 µM), completed with sterile water. We conducted negative controls by replacing cDNA with sterile water and performed standard curves by 5-point dilution ranges with a pool of cDNA (4-fold diluted steps from 0.5 to 2.10^−3^ ng.µL^−1^). We performed a two-steps PCR protocol on a LifeCycler® 480 (Roche Life Science), using the following program: 3 min. of pre-incubation at 95°C, followed by 45 cycles of 15 sec. of incubation (95°C) and 20 sec. of annealing and elongation (60°C) and optical reading. At the end of the program, melting curves were generated (fluorescence optical reading from 65 to 95°C every 0.5 °C) to check amplicon specificity. We normalized expression levels of miRNAs compared to the 5S rRNA as housekeeping gene, which remains stable under experimental conditions. This non-coding small RNA is amplified with forward primer sequence AAGTCCTCGTGTTGCATCCCT. We estimated log_2_ Fold Changes (FC), calculating the relative ratio corrected by PCR efficiency (E) of amplicons (calculated with standard curves: E = 10^(−1 / slope)^) with the following formula : R = (E_miRNA_) ΔCp_miRNA_ ^(control – treatment)^ / (E_5S rRNA_) ΔCp_5S rRNA_ ^(control–treatment)^ (Pfaffl, 2001). We performed statistical analyses by Wilcoxon signed rank tests and considered log_2_ FC significantly different from zero at p.value < 0.05.

### 4. Identification of phenanthrene-responsive miRNAs and their target-genes in *Spartina*

In order to characterize miRNA-regulated gene network in response to phenanthrene, we retained DE miRNAs within *Spartina* species (intraspecific comparisons of control *vs.* phe-treated libraries). We also compared miRNAs expression levels between species from divergent tolerance abilities under phenanthrene-induced stress (interspecific comparisons) (Cavé-Radet *et al*., 2019b). Micro-RNAs expression levels in *S. x townsendii* and *S. anglica* were also compared to those of MPV under phenanthrene-induced stress to identify phenanthrene-responsive miRNAs candidates whose expression was affected by interspecific hybridization and/or genome doubling. Moreover, DE miRNAs under phenanthrene-induced stress between the maternal parent (which exhibits xenobiotic resilience following the *Deepwater Horizon* oil spill; see Wu *et al*., 2012; Robertson *et al*., 2017; Alvarez *et al*., 2018) and the paternal parent were retained.

Annotated targets of candidate phenanthrene-responsive miRNA retained in *Spartina sp.* were compared with *Arabidopsis* (TAIR10 release) gene families potentially involved in xenobiotic detoxification (Edwards *et al*., 2011), or annotated as antioxidants enzymes and transcription factors (TFs). This gene set was composed of 1151 locus we recovered from TAIR gene families index (https://www.arabidopsis.org/browse/genefamily/index.jsp). It includes 94 NAC TFs, 17 Squamosa Promoter binding protein-Like (*SPLs*), 191 α/β hydrolases, 130 ATP-binding cassette (ABC) transporters, 245 cytochromes P450 (*CYPs*), 319 glycosyltransferases (*GTs*), 53 glutathione-S-transferases (*GSTs*), 7 related malonyl-transferases, 3 catalases (*CAT*), 7 superoxide dismutases (*CSD*s, *MSDs* and *FSDs*), 73 class III peroxidases, 8 ascorbate peroxidases (*APX*), and 4 ATP synthase proteins (*ATPS*) (see Supplemental Table S2). These genes are xenome candidates, and are potentially involved in oxidative stress responses induced by xenobiotics in plant tissues, or indirect positive or negative stress tolerance regulators (Singh, 2002; Edwards *et al*., 2005, 2011; Alkio *et al*., 2005; Weisman *et al*., 2010; Dumas *et al*., 2016). Using custom python scripts, we retained phenanthrene-responsive miRNAs involved in the regulation of *Spartina* target genes annotated among these putative stress tolerance regulators.

### 5. Selection of homozygous lines of *Arabidopsis* T-DNA insertional *MIR* mutants

In order to perform a functional validation of phenanthrene-responsive miRNAs regulating transcripts annotated as putative stress tolerance regulators, we first identified, from the set of *Spartina* responsive miRNAs, homologous *MIR* genes in *Arabidopsis* and selected corresponding T-DNA mutant lines. *Arabidopsis thaliana* T-DNA insertional mutants were obtained from the SALK, GABI-KAT and SAIL collections (Colombia (Col-0) genetic background), and from the FLAG collection (Wassilewskija (WS) background) (see Supplemental Table S3). Homozygous plants were selected as described by O’Malley *et al*. (2015). For each mutant, 16 to 20 individual plants were screened. Seed progenies of each mutant were grown on soil in standard conditions and two weeks old rosette leaves were harvested (3-5 mg) and used for genomic DNA extraction as described by Edwards *et al*. (1991). A Two-step genotyping assay was used to identify T-DNA inserts homozygous plants from segregating individuals. A first PCR reaction is performed to confirm that the candidate homozygous line contains a T-DNA insert at the predicted chromosomal location. Primers used were designed by SIGnAL (http://signal.salk.edu/tdnaprimers.2.html) to amplify flanking regions of the insertion site (LP + RP; see Supplemental Table S3). A second PCR reaction uses a universal specific primer (left border LB primer) that spans the predicted T-DNA insertion site. This PCR reaction selectively amplifies the T-DNA/genomic DNA junction sequence (LB + RP; see Supplemental Tables S3 & S4).

### 6. Micro-RNA T-DNA mutants: *in vitro* culture and phenotyping

Homozygous insertional T-DNA mutants were compared with their genetic background under moderate phenanthrene induced stress (50 µM) or high phenanthrene concentration (200 µM), and under control condition in presence of the same volume of ethanol used for solubilization. Seeds were surface-sterilized and sown in Petri dishes on half-strength Murashige and Skoog (MS/2) solid medium containing 0.8 % (w/v) agar-agar type E (Sigma Aldrich) and 1% sucrose. Seeds were germinated vertically in a growth chamber (16:8h light:dark cycle, 8500 lux, 22 °C, 70 % hygrometry) after cold treatment for 48h at 4°C. Within Petri dishes, 10 homozygous seeds per T-DNA mutant line were cultivated under control (0µM) and phenanthrene-induced stress (50 and 200 µM) in technical triplicates. Such phenanthrene concentrations were retained as moderate stress condition (50µM phe) and limiting tolerance concentration (200 µM phe) previously reported in *Arabidopsis* (Dumas *et al*., 2016). Plant phenotypes and photographs were recorded after 7, 14 and 20 days of growth. We measured primary root length as plant tolerance trait to xenobiotic using the ImageJ software (v1.51j8; https://imagej.nih.gov/ij/). At least 10 measures per line and condition were used for statistical analyses. We performed pairwise comparisons between treatments and T-DNA mutants against their related genetic background ecotype using non-parametric Kruskal-Wallis tests (Bonferroni adjusted p.value < 0.05) in R 3.5 (R Core Team, 2015).

### 7. Cloning strategy of miRNAs and target sites

The plasmids used in this study (Supplemental Figure S1) were published by Liu & Axtell (2015) and improved by Moyle *et al*. (2017). For Transient assay systems to validate computationally predicted targets of miRNAs, we used the plasmid pGrDL_SPb which contains the renilla LUC expression cassette acting as internal control to standardize expression between replicates, while the firefly LUC expression cassette containing the predicted target sequence cloned between *Sal*I and *Pst*I is used to report miRNA interaction. Expression plasmid pKENminusbar with multiple cloning sites was used to produce miRNA precursor insertion. Target site adaptors were made by annealing two complementary primers (Supplemental Table S5), synthesized with 5’-phosphate additions and *Sal*I and *Pst*I overhangs (synthesized by Integrated DNA Technologies). A volume of 10 nmol of each primer were mixed together in 60 mL of DNAse-free water, heated to 90°C for 5 min. in a water bath and allowed to slowly cool down to room temperature (at a rate of 1 degree per minute). The resulting adaptors were diluted 100-fold prior to directional cloning into a *Sal*I and *Pst*I digested and Antarctic Phosphatase (New England Biolabs) treated pGrDL_SPb plasmid, and subsequently verified by Sanger sequencing (Australian Genome Research Facility).

The miRNA precursors were synthesized with flanking restriction enzyme sites (GenScript) and cloned between the double Cauliflower mosaic virus (CaMV) 35S promoter and 35S terminator of the expression plasmid pKENminusbar. A similar sized construct, derived from *Capsicum chlorosis* virus, was cloned into pKENminusbar and used as a control (see Moyle *et al*., 2017). This control sequence is predicted to form a secondary structure similarly to a miRNA precursor but does not contain enough homology to produce small RNA capable of interacting with the reporter gene target sequence.

### 8. Agroinfiltration of *Nicotiana benthamiana* leaves

Agroinfiltration was carried out as described by Moyle *et al*. (2017) with minor modifications. Expression plasmids were transformed by electroporation into *A. tumefaciens* strain GV3101 (harboring pSOUP) and plated on LB agar media with rifampicin (25 mg.mL^−1^), kanamycin (50 mg.mL^−1^), and tetracycline (10 mg.mL^−1^) selection. Starter cultures were prepared by inoculating a single colony in 2 mL of LB with rifampicin, kanamycin and tetracycline selection, and grown overnight at 28°C in a shaking incubator. Starter culture was used to inoculate the working culture (20–50 mL), which was grown overnight under the same media, selection and incubation conditions. Working cultures were harvested by centrifugation at room temperature and 2,400g for 15 min. Cell pellets were resuspended in 10 mM MgCl_2_ and the OD600 adjusted to 0.5 prior to the addition of acetosyringone (200 mM). The resulting cell cultures were stored in the dark at room temperature for 4h prior to infiltration. *Nicotiana benthamiana* plants were grown at room temperature and 16h light per day, under LumiBar LED strip lighting (LumiGrow). Seedlings were grown for approximately 3–4 weeks prior to agroinfiltration. Three expanded leaves per plant were infiltrated by applying pressure on the abaxial surface of the leaf with a disposable 5 mL syringe containing the *Agrobacterium* suspension. Each leaf was treated as a biological replicate. Agroinfiltrated plants were incubated for a specified period (typically 72h) in a growth chamber set to 22°C with 16h light per day. Agroinfiltrated leaves were harvested individually, snap-frozen in liquid nitrogen, powdered using a ball mill tissue-lyser (Retsch), and stored at -80°C prior to measurement in dual LUC assays.

### 9. Dual Luciferase assays

Dual luciferase (LUC) assay extracts were prepared using the Dual Luciferase Reporter Assay System kit (Promega). Five mg of powdered tissue was mixed with 100 mL of the passive lysis buffer (PLB) provided in the Dual Luciferase Reporter Assay System kit, and the cellular debris pelleted by centrifugation at 7,500g for 1 min. The supernatant was diluted 20-fold in PLB and 15 µL loaded into a well of a white flat bottom Costar 96 well plate (Corning). The assay was performed using a GloMax 96 microplate luminometer (Promega). The dual injectors were used to introduce 75 mL of luciferase assay reagent and Stop & Glo reagent, respectively, per well. Luciferase assay reagent and Stop & Glo reagent components are provided in the Dual Luciferase Reporter Assay System kit (Promega). Statistical analysis of the resulting data was performed using GraphPad Prism 6 software.

## RESULTS

### 1. Preprocessing of small RNA-Seq and RNA-Seq data: miRNA annotations and mRNA target genes selection

The parental species *S. alterniflora* and *S. maritima*, their F1 hybrid *S. x townsendii* and the allopolyploid *S. anglica* plants were subjected either to phenanthrene treatment or grown under control conditions, as described in the material and methods section. After RNA extraction and sequencing, small RNA-Seq and data preprocessing were performed as previously described in Cavé-Radet *et al*. (2019a). From all combined libraries we obtained 460,502,187 small RNAs (sRNAs) reads. After preprocessing and redundancy removal, we obtained 31,611,711 putative unique sRNA sequences. This procedure consisted of filtering out: (1) the reads with a length lower than 18 bp, over 28 bp, and with more than 2 unknown bases, (2) other non-coding RNA sequences (by Rfam mapping), (3) putative repeat-associated sRNAs; and then collapsing identical reads. Counts of sRNA sequences retained within species and treatments are provided in Table 1. In total, we annotated 594 miRNAs in *Spartina*, distributed between 113 conserved and 481 lineage-specific miRNAs (Cavé-Radet *et al*., 2019a).

**Table 1.** Summary of small RNA-seq data processing. Number of reads retained on each *Spartina* species for control and phenanthrene-treated libraries are presented.

| Species | *S. alterniflora* |  | *S. maritima* |  | *S. x townsendii* |  | *S. anglica* |  |
| --- | --- | --- | --- | --- | --- | --- | --- | --- |
| Treatment | Control | PHE | Control | PHE | Control | PHE | Control | PHE |
| Raw reads | 37551835 | 60444313 | 17828306 | 32274148 | 100752127 | 125380090 | 46650696 | 39620672 |
| Clean reads | 22231291 | 31443611 | 11042707 | 21336208 | 73548538 | 97497156 | 35270284 | 25825213 |
| Putative sRNA reads | 5718925 | 7367871 | 3389799 | 4676878 | 25809487 | 52366512 | 20524897 | 9612503 |
| Putative sRNA reads (non redundant) | 2252788 | 2284748 | 1613905 | 1853687 | 7238893 | 9208854 | 4434910 | 2723926 |

From RNA-Seq, we obtained about 300 million of paired reads in total, all libraries combined. Sequencing quality checking (Phred Q-score > 30) was performed using the FastQC software (Andrews, 2010) prior to mapping reads on contigs selected from putative miRNA target genes (Cavé-Radet *et al*., 2019a) and *Spartina* transcripts expressed under phenanthrene-induced stress conditions.

### 2. Effects of hybridization and polyploidy on miRNAs expression under phenanthrene induced stress

Several publications pointed out that miRNAs may arise from transposable elements and DNA rearrangement following genomic ‘shock’. Hence, we first investigated if hybridization and genome doubling induced *MIR* expression repatterning, specifically activated under xenobiotic induced stress. To this end, we compared miRNAs distribution between the parental species, *S. alterniflora* and *S. maritima*, the F1 hybrid *S. townsendii*, and the allopolyploid *S. anglica* (figure 1A, B) under stressed conditions. No species-specific conserved miRNAs were identified, as most miRNAs were expressed in all four species. This was also observed in novel miRNAs sets, but to a lesser extent, as only very few novel miRNAs were found to be species-specific (4, 4, 2 and 2 miRNAs in each of the parents, the F1 hybrid and the polyploid, respectively). Among 481 *Spartina*-lineage specific miRNAs, 94.2% of them are common. Meanwhile, it is interesting to notice that 6.65% of novel miRNAs emerged following hybridization and genome doubling in these conditions. These results indicated that hybridization and genome doubling do not induce dramatic changes in the emergence of *MIR* genes under xenobiotic induced stress. However, the global level of miRNAs expression in the parents and the MPV compared to *S. townsendii* and *S. anglica* showed very contrasted level between control and phenanthrene treated conditions after hybridization and genome doubling (data not shown). All together, these data indicated that hybridization and genome doubling mostly affect miRNAs expression levels rather than the emergence of newly expressed *MIR* genes. We performed experimental validation (RT-qPCR) for *in silico* expression profiles of 11 miRNAs (Supplemental Figure S2). These miRNAs were selected following primer design and we retained miRNAs with the best primer parameters for PCR amplification and specificity (Supplemental Table S1). Due to very low amplification efficiency for Spar-new-miR_236 (E = 1.508), we removed this latter from the analysis. Overall, we found the same tendency of the RT-qPCR and small RNA-seq data.

**Figure 1.**
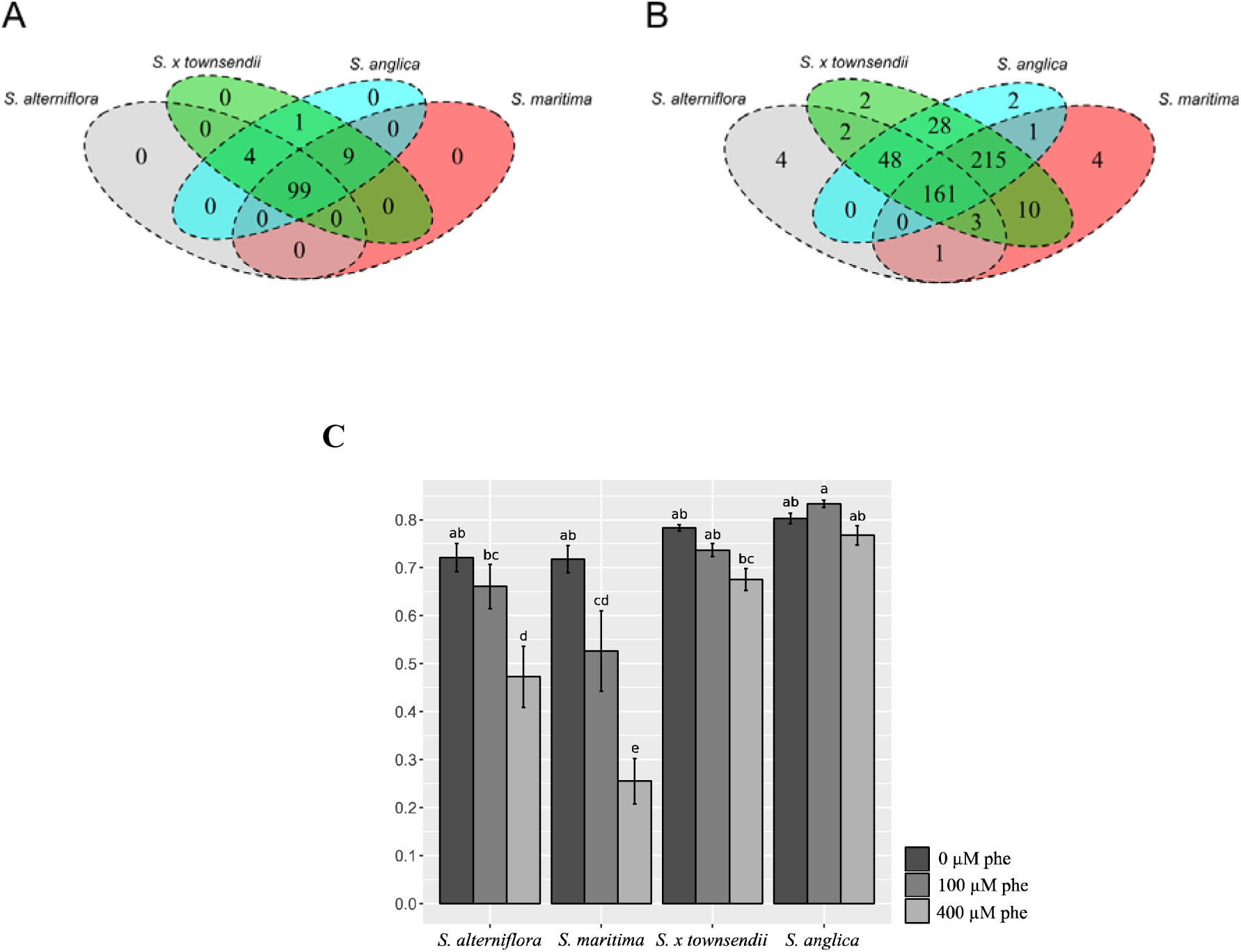
Venn Diagrams of conserved (A), lineage-specific (B) *Spartina* miRNAs expressed under phenanthrene-induced stress and *Fv/Fm* ratios (C). The *Fv/Fm* was used as an index of the photosystem II activity in response to stress. Low tolerance abilities were reported in *S. maritima* and *S. alterniflora*, as the *Fv/Fm* ratios was significantly reduced since 100 µM phe treatments. In contrast, hybrid species *S. x townsendii* and *S. anglica* present enhanced tolerance abilities compared to parental species. Values annotated with different letters are significantly different according to *post hoc* Tuckey HSD test following 2-way ANOVA (p.value < 0.05).

Our previous published data (Cavé-Radet *et al*., 2018b), and additional comparative physiological analysis (figure 1 C) showed enhanced tolerance to phenanthrene in the F1 hybrid *S. x townsendii*, which may be attributed to heterosis effect, and in the allopolyploid *S. anglica*, when compared to parental species. According to such contrasting tolerance levels between species, we reasoned that the hybrid and the polyploid intraspecific DE miRNAs, and interspecific miRNAs following hybridization and polyploidization under phenanthrene treatment may play a pivotal role in xenobiotic tolerance. Hence, we performed differential expression analysis of miRNAs under phenanthrene induced stress by comparing parental species *S. alterniflora*, *S. maritima*, the F1 hybrid *S. x townsendii* and the allopolyploid *S. anglica* expression levels in phe-treated *vs.* control conditions. In total, we identified 103 significantly differentially expressed (DE) phenanthrene responsive miRNAs: 20 in *S. alterniflora*, 14 in *S. maritima*, 54 in *S. x townsendii*, and 33 in *S. anglica*. A close analysis of miRNAs dynamic in control and stressed condition revealed that significant changes in miRNAs expression occurred, indicating that miRNAs transcription was impacted by hybridization, polyploidy and genome doubling *per se.* Under phenanthrene induced stress, most miRNA transcriptional changes were detected following hybridization (12.87%) compared to genome doubling (4.84%) or between the MPV against *S. anglica* (5.62%). Transcriptional changes in miRNA expression following allopolyploidization, and between the parental species (19.19%) are presented in figure 2.

**Figure 2.**
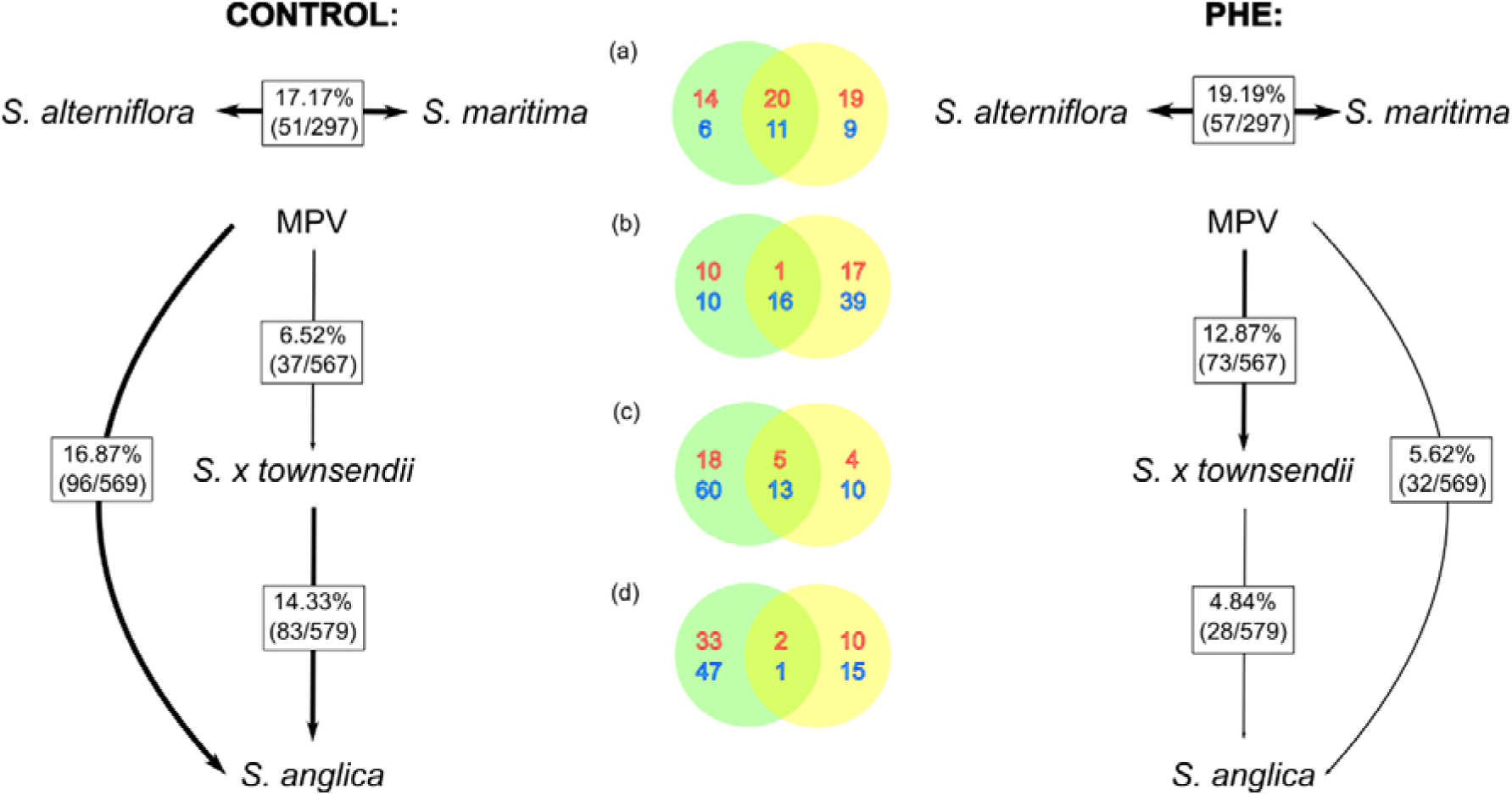
Transcriptional changes of miRNAs expression between parental species (a) species following hybridization (b), allopolyploidy (c) and genome doubling *per se* (d) under phenanthrene-induced stress (PHE) compared to control condition (Cavé-Radet *et al*., 2019a). The percentages are indicating the proportions of expression changes, Venn diagrams are representing the number of up-regulated miRNAs (in red) and down-regulated (in blue) between control (green circles) and phenanthrene-induced stress conditions (yellow circles). Between parental species, up- or down-regulated miRNAs are referred comparing *S. alterniflora* to *S. maritima*.

Results from intraspecific differential analysis (control vs PHE-treated) are presented on a heatmap (figure 3). We found that hierarchical clustering based on expression profiles among *Spartina* species did not bring out any obvious miRNA clusters. Nonetheless, most of DE miRNAs have been identified within species, subsequently some of them such as Spar-miR395d, Spar-new-miR_384, Spar-miR156c and Spar-miR157 are significantly up-regulated in both *S. x townsendii* and *S. anglica*. For other miRNAs, significantly opposite expression profiles are observed between species, for example Spar-new-miR_154 or Spar-miR160 were up-regulated and down-regulated respectively in the F1 hybrid and the allopolyploid. Major DE miRNAs (log_2_ FC > 5 or < -5), which represent 27 sequences are listed in Supplemental Table S6.

**Figure 3.**
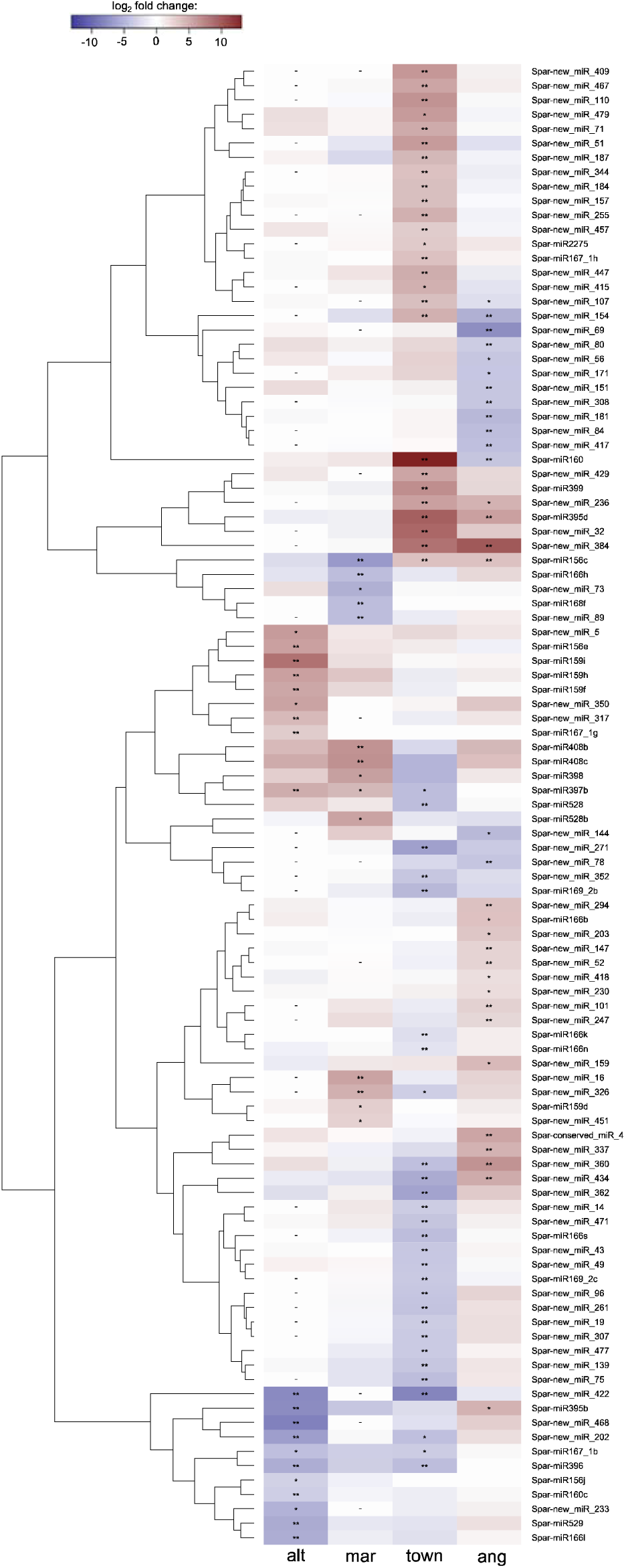
Clustering and heatmap of 103 differentially expressed miRNAs in response to phenanthrene (alt: *S. alterniflora*; mar: *S. maritima*; town: *S. x townsendii*; ang: *S. anglica*). DE Spar-miRNAs were clustered according to their expression patterns between species (*: p.value < 0.1; **: p.value < 0.01; -: miRNA not expressed in a given species).

In complement, we analyzed miRNAs expression changes under phenanthrene in response to hybridization, polyploidization and genome doubling *per se* by comparing *S. x townsendii* and *S. anglica* to the MPV and *S. anglica* to *S. x townsendii* respectively (figure 4). Nonadditive expression profiles were detected through 73 DE miRNAs following hybridization (18 up-regulated and 55 down-regulated), by comparing *S. anglica* to the MPV, we identified 32 DE miRNAs (9 up-regulated and 23 down-regulated) and 28 DE miRNAs following genome doubling (12 up-regulated and 16 down-regulated).

**Figure 4.**
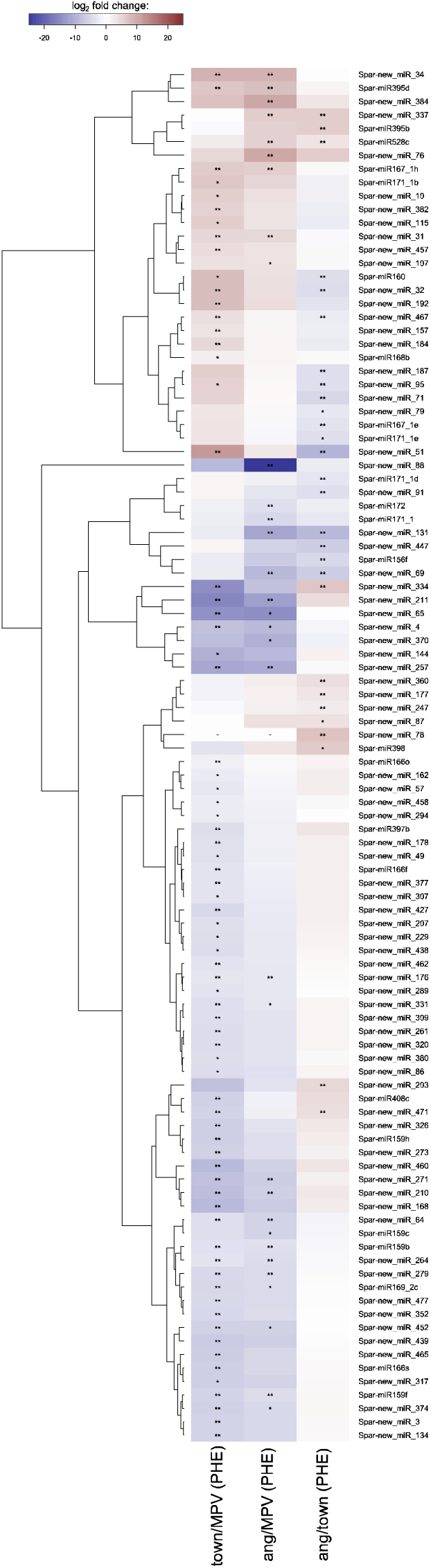
Clustering and heatmap of differentially expressed miRNAs in response to hybridization (*S. x townsendii* compared to MPV), genome doubling *per se* (*S. anglica* compared to *S. x townsendii*) and between *S. anglica* compared to MPV under phenanthrene-induced stress (town: *S. x townsendii*; ang: *S. anglica*; MPV: Mid-Parent Value). DE miRNAs are clustered according to their expression patterns across comparisons (*: p.value < 0.1; **: p.value < 0.01; -: miRNA not expressed in species).

### 3. Identification of differentially expressed transcripts among *Spartina* species in response to phenanthrene

Using RNA-Seq data, we studied the expression of candidate miRNA target genes potentially involved in phenanthrene responses and performed the same comparisons as described for miRNAs. This procedure aimed at identifying stress-responsive transcripts, and to compare miRNA expression levels to those of their putative target genes. In total, we followed the expression of 9,714 contigs based on the counting matrix we obtained from mapping RNA-Seq reads. Only transcripts with sufficient mapping depth (10 reads minimum in average among libraries) which represent 3,267 contigs were processed by EDAseq (Risso *et al*., 2011) and DESeq2 (Love *et al*., 2014) for differential expression analysis. We identified 36 DE transcripts in *S. alterniflora* in response to phenanthrene, 3 in *S. maritima*, 13 in *S. x townsendii*, and 22 in *S. anglica*. From intraspecific comparisons in response to phenanthrene, low coverage in DE contigs between species was detected (Figure 5A), and most of contigs (92%) appeared DE in one or other of species suggesting the occurrence of species-specific molecular mechanisms in response to stress.

**Figure 5.**
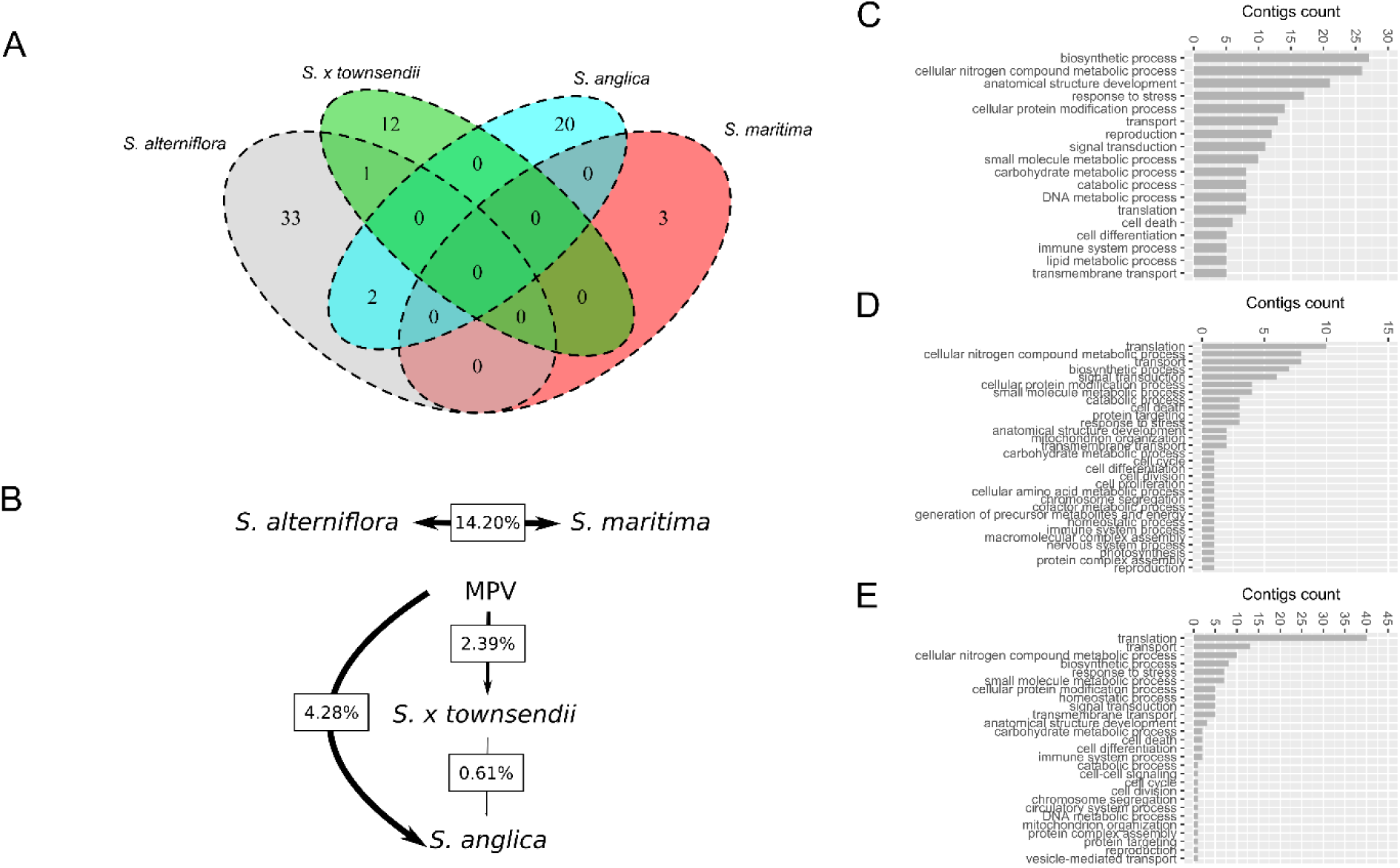
DE target genes (contigs) identified between *Spartina* species in response to stress (A). Transcriptional changes in response to stress between parental species, hybridization (*S. x townsendii* compared to MPV), genome doubling *per se* (*S. anglica* compared to *S. x townsendii*) and between *S. anglica* compared to MPV are presented as percentages (B). Biological process GO terms distribution associated to DE contigs followed by RNA-Seq between parental species (C), following hybridization (D), and allopolyploidization (E).

Transcriptional changes detected between the parental species, and following allopolyploidization under phenanthrene are presented Figure 5B. As observed for miRNAs, most of transcriptional changes were observed between parental species (14.20%). In *S. x townsendii*, we observed 2.39% transcriptional changes, and 4.28% comparing *S. anglica* to the MPV, against only 0.61% between *S. anglica* and *S. x townsendii*. Biological process GO terms related to DE contigs identified among species in response to stress are detailed on Figure 5 between parental species (C), DE contigs between *S. x townsendii* (D) and *S. anglica* (E) compared to the MPV. Most of differentially expressed contigs between the parental species span a myriad of functions annotated among ‘biosynthetic process’, ‘cellular nitrogen compounds metabolic process’, ‘anatomical structure development’ or ‘response to stress’, which could reflect the evolutionary divergence between the parental species *S. alterniflora* and *S. maritima*. Contrary to biological GO terms related to DE contigs following hybridization and polyploidy, only few differentially represented biological function are represented, interestingly the most represented was ‘translation’, which might be related to heterosis and/or genome doubling effects.

### 4. Many D1E miRNAs targeting xenome genes under phenanthrene induced stress are reprogrammed following hybridization and genome doubling

We examined to what extent DE miRNAs targeting the xenome components are affected following hybridization and polyploidy under phenanthrene induced stress. Among phenanthrene responsive miRNAs we retained those targeting genes annotated as putative xenome gene candidates. Interestingly, some phenanthrene responsive miRNAs are up-regulated in the hybrid and the polyploid species such as Spar-new-miR_3 and the *MIR*156 family (Spar-miR156c, Spar-miR156d, Spar-miR156e and Spar-miR156h), these miRNAs target *SPL* homologous genes. Indeed, down-regulation of *SPLs* has been correlated with enhanced growth, development, and tolerance to several stress. Moreover, we detected down-regulation of several novel miRNAs Spar-new-miR_1, Spar-new-miR_426 and Spar-new-miR_151 and the conserved *MIR*159 family (Spar-miR159d and Spar-miR159j), these miRNAs are targeting *GTs* and *CYPs.* Other down-regulated novel miRNAs (Spar-new-miR_56, Spar-new-miR_72, Spar-new-miR_181, Spar-new-miR_447 and Spar-new-miR_316) targeting *CYPs* and α/β hydrolases were identified. Consequently, up-regulation of all these xenome genes may impact degradation and tolerance abilities to oxidative stress generated by xenobiotic in plant tissues.

### 5. Heterologous functional analysis of phenanthrene-responsive conserved miRNAs

In order to identify the most determinant miRNAs involved in xenobiotic tolerance, we first browsed *Brachypodium* T-DNA mutant collections. However, only very few *MIR* gene - mutants were found. Consequently, we focused our research on *Arabidospsis* T-DNA mutants focusing on knock-out phylogenetically-conserved miRNAs. Hence, we collected 39 T-DNA insertional *Arabidopsis* mutants of DE putative phenanthrene responsive miRNAs (all conserved *Spartina* miRNAs; see supplemental Table S3), mainly in the hybrid and the allopolyploid, and resulting from the hybridization or the allopolyploidization (differentially expressed in the F1 hybrid or the allopolyploid compared to the MPV). Genotypes of each line were first screened, based on the T-DNA insertion in the genome, to identify homozygous individual plants for each miRNA mutant, and only seeds of homozygous individual plants of each line were screened. Interestingly, among the most contrasted phenotype under phenanthrene induced stress, the homozygous line FLAG_090F02 harboring a T-DNA insertion in an ath-*MIR*156 (AT5G11977) was found to be sensitive to phenanthrene (Figure 6). When cultivated in low xenobiotic concentration (50 µM phe), this mutant showed no difference in root growth but senescent phenotypes were observed in aged leaves, which turned yellow and lost all the chlorophyll in comparison to wild type (Figure 6B). This sensitive phenotype was more contrasted under high phenanthrene concentration (200 µM), as the first leaves developed very slowly and primary roots showed growth inhibition (Figure 6C). We then phenotyped another independent allelic homozygous mutant FLAG_104E09, where the T-DNA was inserted in the exon of AT5G11977, this later showed the same tendency under high phenanthrene concentrations (Figure 6D).

**Figure 6:**
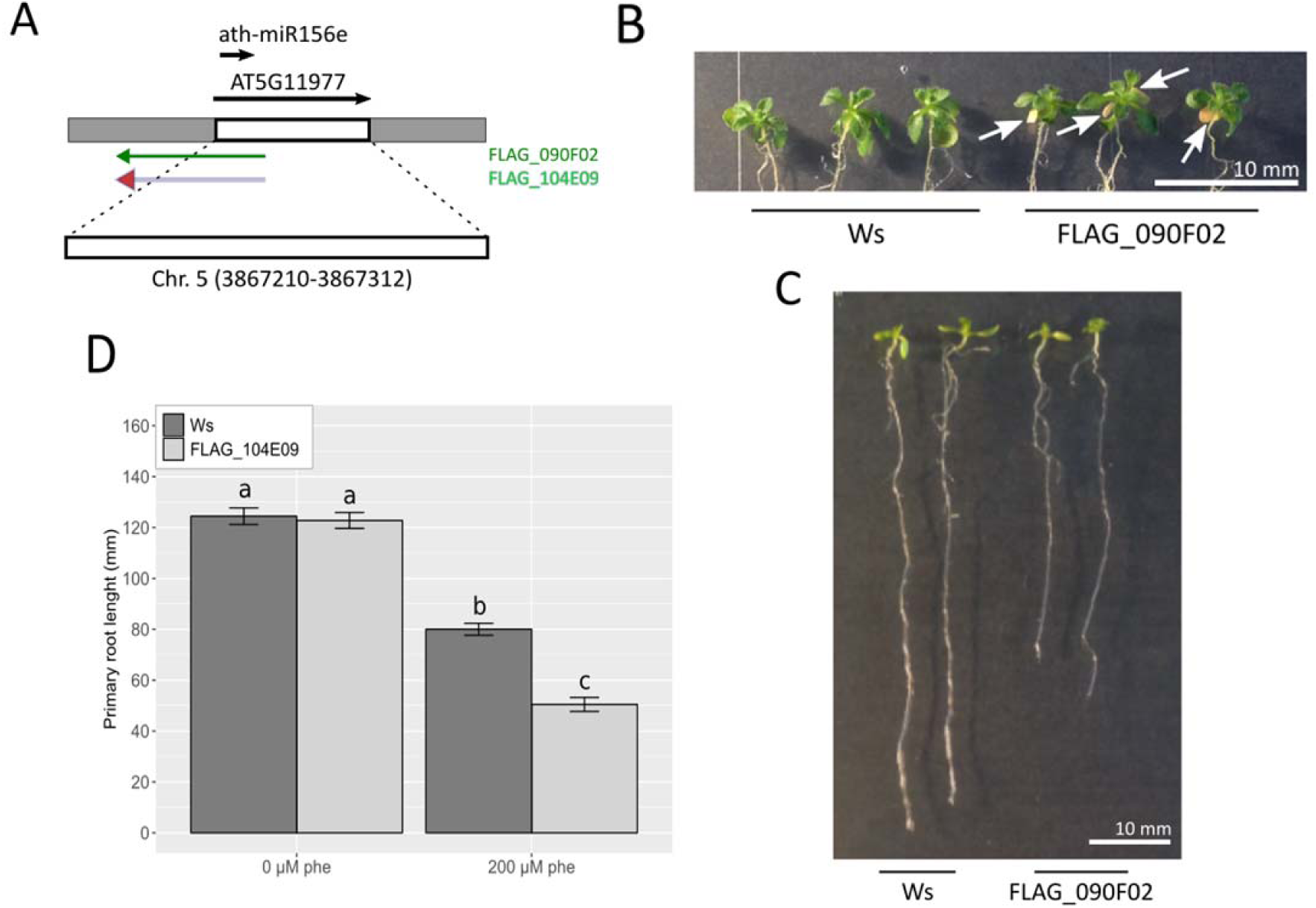
Functional analysis of sensitive ath-MIR156e T-DNA mutant following phenanthrene treatments. Insertions of FLAG_090F02 and FLAG_104E09 T-DNA lines in the *Arabidopsis* genome (inserted in exon of *MIR* gene AT5G11977) are presented (A). Homozygous mutant line FLAG_090F02 showed sensitive phenotype as aged leaves turned yellow and lost all the chlorophyll after 14 days at low phenanthrene containing medium (50 µM) (B). When cultivated in higher phenanthrene concentration containing medium (7 days on 200 µM), root growth of FLAG_090F02 was reduced compared to Ws ecotype. Representative results are shown (C). Primary root lengths (± ES) of Allelic homozygous FLAG_104E09 mutant line, and the related genetic background grown under high phenanthrene content (20 days on 200 µM) were compared and plotted (D). This allelic mutant showed the same tendency. Bars annotated with different letters are significantly different according to pairwise Kruskal-Wallis tests (Bonferroni adjusted p.value < 0.05).

In contrast, homozygous SALKseq_071642 (ath-*MIR*156 family homolog, see Figure 7C) and SAIL_770_G05 (ath-*MIR*159b) T-DNA lines exhibited tolerant phenotypes under 200 µM phenanthrene induced stress (Figure 7E). Here, while the corresponding background genotype Col-0 showed growth inhibition between 30 to 40%, the SALKseq_071642 did not show any significant inhibition (Figure 7C). Concerning SAIL_770_G05 mutant line, the root growth decreased of 12.5% under phenanthrene induced stress, significantly different to the corresponding background genotype Col-0 which decreased of 32% in primary root length in response to stress (Figure 7E).

**Figure 7.**
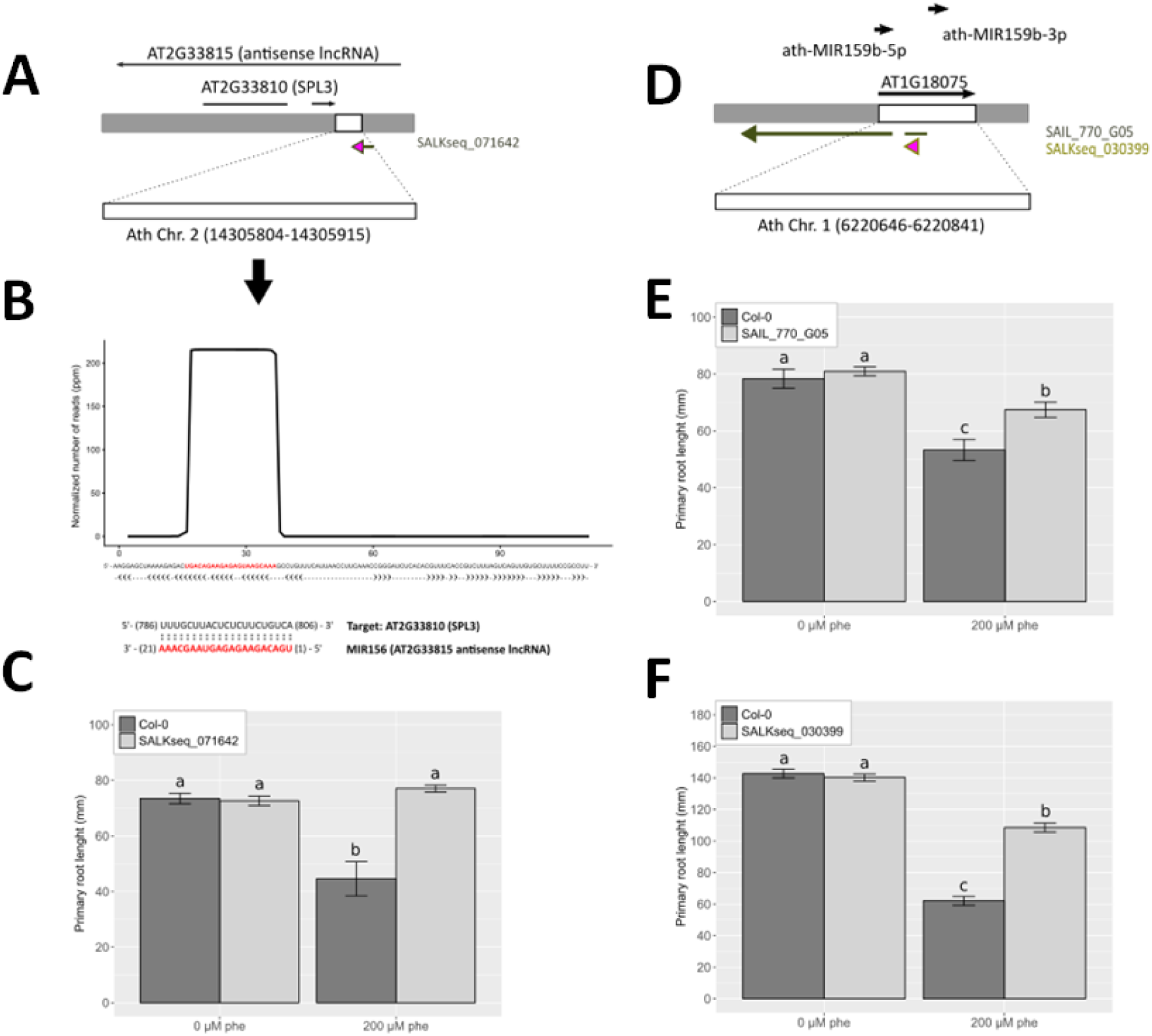
Functional analysis of the most tolerant T-DNA mutants. Insertion profiles of SALKseq_071642 (A) and SAIL_770_G05 (D) T-DNA lines in Arabidopsis genome (respectively inserted in 300-UTR3 of the predicted MIR156 and in exon of *MIR159b* gene AT1G18075). Putative novel MIR156 miRNA identified in *Arabidopsis* lncRNA AT2G33815 (B). *Spartina* miRNAs were mapped on *Arabidopsis* genome (Chr. 2 positions 14305804 to 14305915), which present stem-loop precursor profile (B). This putative novel-MIR156 sequence (in red) was predicted to target *SPL3* by perfect matching with the opposite complementary strand (B). After 14 days cultivated under 0 and 200 µM phe, primary root lengths (± ES) of T-DNA mutants and their related genetic background were compared and plotted (C and E). Functional analysis of the allelic mutant SALKseq_030399.2 showed the same tendency as the SAIL_770_G05 line (F). Bars annotated with different letters are significantly different according to pairwise Kruskal-Wallis tests (Bonferroni adjusted p.value < 0.05).

The T-DNA insertions in the FLAG_090F02 and SAIL_770_G05 lines were found in exons of ath-*MIR*156e (AT5G11977) and ath-*MIR*159b (AT1G18075) well annotated miRNA primary transcripts (Figure 6A and 7D). However, in SALKseq_071642 mutants the T-DNA insertion was found in the exon of an antisense long non-coding RNA (lncRNA; AT2G33815), which overlaps with *SPL3* (inserted in 300-UTR3 of AT2G33810; Figure 7A). A closer analysis of this lncRNA sequence revealed a typical miRNA precursor with characteristic hairpin structure located between positions 14305804 and 14305915 on Chr2. *In silico* analysis of this sequence, revealed a high and confident homology on this putative miRNA precursor (predicted miR156 candidate) from mapping *Spartina* deep sequencing reads (average mapping depth up to 200 RPM; see Figure 7B) on *Arabidopsis* genome, and analysis of the secondary structure of the flanking sequence. Hence, we hypothesized that the annotated lncRNA encoded a functional miRNA from the *MIR*156 family, which potentially target *SPL3* (as shown in Figure 7B).

### 6. Experimental validation of ath-*MIR*159b, ath-*MIR*156e, and the predicted-miR156 interactions and their putative targets

We extend miRNA/target gene validation from merely *in silico* and descriptive analysis to functional studies. For target genes, we selected sequences with the better prediction ranking scores (≤ 3.0 from psRNATarget; Dai & Zhao, 2011) for miRNAs ath-*MIR*159b, ath-*MIR*156e, and the predicted *MIR*156 candidate (lncRNA, AT2G33815).

The dual LUC assays revealed significant repression of the firefly LUC reporter when ath*-MIR*159b and the predicted target sequence of the *MYB33* domain protein (AT5G06100) were co-agroinfiltrated (Figure 8). In parallel, ath-MIR156e induced significant inhibition in the presence of *SPL2* and *SPL4*, but no difference in expression level was observed when *SPL13B* target sequences was co-infiltrated with the ath-MIR156e precursor or the control precursor (Figure 8). This may be explained by low complementarity between the indicated miRNA and the *SPL13B* target sequence. Consequently, we hypothesized that ath-MIR156e does not interact with all *SPL* genes but only with specific ones to control tolerance to xenobiotic organic pollutants. Surprisingly, no difference in expression level was observed when *SPL3* target sequence was co-infiltrated with the predicted MIR156 candidate (lncRNA) precursor compared to internal control, even if perfect complementarity between miRNA and its target was reported. Moreover, we did not find any repression of the firefly LUC reporter gene when the predicted MIR156 candidate was co-infiltrated with *SPL4* and *SPL13B* predicted target sequences (Figure 8). These results indicated that even the lncRNA (AT2G33815) meets some important criteria to be classified as miRNA precursor, it may not be able to produce mature and functional miRNA in *Arabidopsis*.

**Figure 8.**
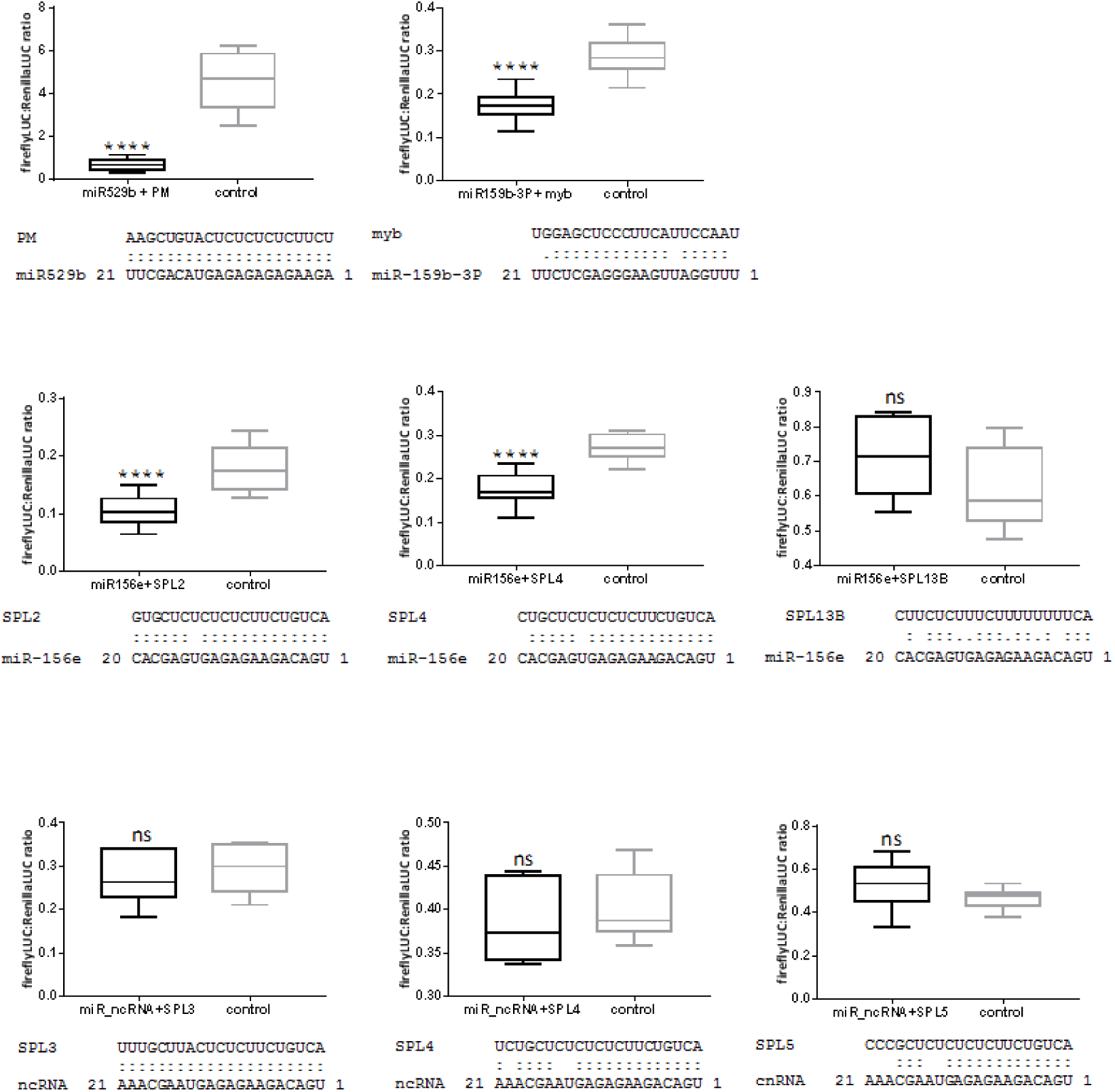
Experimental validation of computational interaction of *Arabidopsis* miRNAs (homologous to phenanthrene-responsive miRNAs in *Spartina*) and their targets. Each control assay consists on the agrobacterium co-infiltration of a non-processed precursor construct culture with the indicated target construct. miR529b-mediated repression of perfectly matching (PM) target reporter gene expression was used as a positive control. The controls consist of assays where the precursor cultures were replaced with a precursor construct culture that does not target the indicated miR PM sequence. Target genes are: MYB33 domain protein (AT5G06100), *SPL2*: Squamosa promoter binding protein-like 2 (AT5g43270), *SPL3*: Squamosa promoter binding protein-like 3 (AT2G33810), *SPL4*: Squamosa promoter binding protein-like 4 (AT1G53160), *SPL5*: Squamosa promoter binding protein-like 5 (AT3G15270), *SPL 13B*: Squamosa promoter binding protein-like 13B (AT5G50670). The ncRNA referred to the MIR156 candidate predicted from the antisense lncRNA (AT2G33815). Data were depicted as box and whisker plots with interquartile range boxes and min/max whiskers (n=6). Asterisk represents statistically significant differences using a simple two-tailed unpaired *t*-test with ****p.value < 0.0001.

## DISCUSSION

In this study, we analyzed stress-responsive miRNAs in polyploid *Spartina* species under phenanthrene, a common organic pollutant from the PAHs. This represents the first miRNA profiling in *Spartina* species under experimental stressful conditions, using the annotation dataset developed from homology searches and *de novo* predictions recently published (Cavé-Radet *et al*., 2019a). Here, we identified phenanthrene-responsive miRNAs and performed DE analyses to highlight the impact of allopolyploidization in reprogramming miRNAs, their target genes, and other transcripts potentially involved in phenanthrene tolerance. To date, only salt-responsive miRNAs were explored in *S. alterniflora*, based on microarray transcriptional profiling (Qin *et al*., 2015) and semi-quantitative expression analysis from sequences identified by homology searches to miRNAs deposited in public databases (Zandkarimi *et al*., 2015). All combined, these studies examined 91 miRNAs, but low overlap with our datasets was identified (only 9 in common). Thus, our dataset (594 miRNAs) provides an increased range of miRNAs potentially related to stress-induced regulations.

### Hybridization and genome doubling reprogrammed miRNA expression under phenanthrene-induced stress in *Spartina*

As presented, most miRNAs transcriptional changes were observed between the parental species (19.19%), consistently with previous results obtained by Chelaifa *et al*. (2010b,a) on mRNAs under standard conditions. This likely reflects evolutionary history of these two sister species that diverged 2-4 Mya (Rousseau-Gueutin *et al*., 2015). In response to allopolyploidization, major miRNAs expression changes (non-additive parental patterns) were detected following hybridization (12.87%) compared to genome duplication *per se* (4.84%), contrasting with recently reported results obtained in standard condition where major changes were observed comparing *S. anglica* to the MPV (16.87%) rather than following hybridization (6.52%, Figure 2; Cavé-Radet *et al*., 2019a). Such changes in miRNAs expression patterns in *S. x townsendii* and *S. anglica* may rely on a buffering effect in response to the genomic “shock” following genome merger and redundancy such as reported in *Arabidopsis* hybrid genomes by Ha *et al*. (2009) or in previous global transcriptome analyses in Spartina (Chelaifa *et al*. (2010b). Here, it appears that DE miRNAs in phe-treated plants following hybridization may impact heterosis, as reported under standard culture experiments in rice (He *et al*., 2010), tomato (Shivaprasad *et al*., 2012), sugarcane (Zanca *et al*., 2010) or wheat (Kenan-Eichler *et al*., 2011). In response to stress, this can lead to increased tolerance abilities in *S. x townsendii* compared to the parental species. Some remarkable miRNAs such as Spar-new-miR_429, Spar-miR399, Spar-new-miR_236, Spar-miR395d, Spar-new-miR_32, Spar-new-miR_384 and Spar-miR156c were particularly interesting. These miRNAs are clustered together on the heatmap Figure 3 and present up-regulation patterns in response to phenanthrene in *S. x townsendii* and *S. anglica*. Such miRNAs may then impact the differences in tolerance abilities observed following allopolyploidization (Cavé-Radet *et al*., 2019b). Recently, we reported from comparative physiological analyses that the combination of hybridization and genome doubling events impact tolerance abilities to PAHs in *Spartina*. Such features are most likely impacting *Spartina* species ecology, like resilience abilities to chemical pollution, or the rapid spread of *S. anglica* in Europe, and following its introductions in North America, China, Australia, where it is now listed among the 100 worst invasive alien species (IUCN, 2000).

### Phenanthrene-responsive miRNAs in *Spartina* and target genes

We detected phenanthrene responsive miRNAs candidates by comparing expression patterns within species (phenanthrene-treated *vs.* control, intraspecific comparisons), combined with interspecific comparisons.

From intraspecific comparisons, 103 DE miRNAs in response to phenanthrene were identified, all species combined. Most of RNA-seq expression profiles were validated by RT-qPCR, and confirming this DE analysis (Supplemental Figure S2). Most of DE miRNAs present significant expression regulation patterns that are species-specific, as low consistence between species were observed based on hierarchical clustering analyses. Surprisingly, we rarely observed opposite miRNAs regulations between species from contrasted tolerance abilities (*e.g.* up- and down-regulated miRNAs in response to phenanthrene between species). Such patterns are principally contrasting *S. x townsendii* to *S. anglica*, as observed for Spar-new-miR_107, Spar-new-miR_154, Spar-miR160, Spar-new-miR_360 and Spar-new-miR_434 (see Figure 3). More specifically, we identified DE miRNAs in *S. anglica* where enhanced tolerance abilities are observed following allopolyploidization (Cavé-Radet *et al*., 2019b). They represent 33 *Spartina* lineage-specific miRNAs (19 up-regulated and 14 down-regulated). Most of them (27) are lineage-specific *Spartina* miRNAs. Moreover, the 27 miRNAs with extreme FC values reported in Supplemental Table S6 are to consider in further studies focusing on the role of miRNAs in response to PAHs. Based on inter-specific comparisons, we retained 164 unique candidate phenanthrene-responsive miRNAs (46 conserved and 118 lineage-specific Spar-miRNAs). Altogether, this provides a set of 220 candidates (63 conserved and 157 lineage-specific Spar-miRNAs) which include reprogrammed miRNAs identified within species in response to stress and between closely related species from contrasted tolerance abilities. These sets of miRNAs may target positive or negative stress tolerance regulators respectively and represent candidates of interest potentially involved in gene regulation pathways in response to stress.

Based on annotations of the putative targeted genes by phenanthrene-responsive miRNAs retained in *Spartina*, we identified 15 miRNAs (5 conserved miRNA from MIR156 and MIR159, and 10 lineage-specific Spar-miRNAs) which target genes potentially involved in xenobiotic detoxification. A recent study published by Li *et al*. (2017) reported 11 stress-responsive miRNAs from wheat watered 2 days with phenanthrene supplemented Hoagland solution (1 mg. L^−1^). In their study, the authors focused on roots to perform miRNAs DE analysis in early xenobiotic exposure (0, 12, 24 and 48h). Our study was performed after 5 days of phenanthrene exposure to specifically focus on detoxification responses, once xenobiotics accumulate in shoot tissues. Among Poaceae, we crossed phenanthrene-responsive miRNAs from *Spartina* and *Triticum* species, but we observed low overlap between the two taxa with only two commonly DE miRNAs. We found Spar-miR156e (UGACAGAAGAGAGUGAGCACA) up-regulated in both taxa, and Spar-miR398 (UGUGUUCUCAGGUCGCCCCCG), up-regulated in *Spartina* leaves but down-regulated in wheat roots. *MIR*156 and *MIR*398 families are shared among spermaphytes, and their impact in stress responses are also well described (Ding *et al*., 2010; Aung *et al*., 2015). In *Spartina*, similar functions may be expected in response to PAH xenobiotics, and the opposite expression patterns we observed compared with wheat could result from spatiotemporal differences (*e.g*. organs, time exposure) between studies and/or from species divergence.

Depending on comparisons performed, we retained in total 7 phenanthrene-responsive miRNAs (Spar-miR156e, Spar-miR156c, Spar-new-miR_181, Spar-miR156d, Spar-miR156h, Spar-new-miR_426 and Spar-new-miR_3) where opposite expression responses compared to their putative target genes were observed (Supplemental Figure S3) even if no significant expression regulations of these target genes were observed (mostly annotated as *SPLs*, and as *GT* or α/β hydrolases). However, the others phenanthrene-responsive miRNAs (Spar-miR159d, Spar-miR159j, Spar-new-miR_1, Spar-new-miR_56, Spar-new-miR_72, Spar-new-miR_151, Spar-new-miR_316 and Spar-new-miR_447) remain powerful candidates potentially involved in RNAi regulation of genes involved in tolerance to xenobiotics. Moreover, DE analyzes performed on *Spartina* contigs (putative miRNA target genes and *Spartina* transcripts assembled under phenanthrene-induced stress) revealed various phenanthrene-responsive genes. As described above, most of them are specifically DE in one or other species and suggest different responses in transcriptomic profiles under phenanthrene, which may be related to enhanced tolerance abilities detected in *S. anglica* following allopolyploidization. These contigs need to be considered in further analyzes focused on molecular responses to PAHs in plants.

### *MIR*159 and *MIR*156 regulatory modules are evolutionary targeted to adapt the xenome expression and developmental plasticity responses under phenanthrene-induced stress

MiRNAs sequencing and annotation of four highly redundant *Spartina* genomes revealed 481 new and 131 conserved miRNAs. Meanwhile, the identification of the most determinant or ‘the master regulator’ miRNAs following hybridization and allopolyploidy involved in xenobiotic tolerance is a challenging task. In the present investigation we decided to use *a* without *a priory* strategy, based on a phenotypical screen of heterologous *Arabidopsis* T-DNA mutants. We conducted functional validation experiments in relatively distant plant lineages (Monocots *vs.* Dicots), and kept focused on conserved *MIR* gene families within embryophytes (Taylor *et al*., 2014). This strategy allowed to identify among 39 independent mutant lines, contrasted phenotype of two miRNAs mutants. Interestingly, the detected DE miRNAs showed that hybridization and genome doubling impacted negatively almost all *MIR*159 genes (see figure 9A). Targets with the highest prediction ranking scores (above 5.0 from psRNATarget; Dai et al. 2018) (see supplementary table S7) (i) were found to code xenome products of almost all xenobiotics detoxification steps as shown in figure 9B, (ii) and also code for many superoxide dismutases, known to be involved in ROS scavenging under xenobiotics induced stress. We hypothesized that *MIR*159s were selected as a negative master family gene that under xenobiotic pressure allowed relaxation of the xenome and ROS scavenging targets genes. On the other hand, MIR159 is an evolutionary conserved miRNA that targets MYB transcription factors to function both as a switch and as a tuning mechanism, modulating the expression of its targets in response to endogenous and exogenous signals (Alonso-Peral *et al*., 2012). MYB transcription factors and miR159 interactions are involved in many stress responses like UV-B or drought (Roy, 2016; Li *et al*., 2016; Chen *et al*., 2018). However, their function in xenobiotic detoxification or tolerance related mechanisms were never described to our knowledge. Hence, the dual LUC assays revealed very significant silencing of the firefly LUC reporter when ath-MIR159 and the predicted target sequence of the MYB33 domain protein were co-agroinfiltrated, indicating that genomic stress ‘shock’ may act throughout miR159/MYB33 module under xenobiotic induced stress. Guo *et al*. (2017) showed that the loss of miR159 resulted in miR156 increased level. MiRNAs from the MIR156 family target transcription factors of *Squamosa promoter binding-like* (*SPL*) family, which are crucial genes involved in plant development (Xu *et al*., 2016; Hyun *et al*., 2017). The repression of miR156 by miR159 is predominantly mediated by MYB33/62, MYB33 simultaneously impacts the transcription of MIR156 genes, as well as their target SPL9 by directly binding to the promoters of these genes. Interestingly, the STTM159 transgenic resulted in the increased expression of target gene MYBs, and exhibited short stature along with smaller organ size, highlighting the importance of MIR159 genes in promoting cell division and organs development (Zhao et al. 2017). In the present work, functional analysis of knock-out mutant harboring a T-DNA insertion in the exon of miR156 homolog showed sensitive phenotype. These data fit well with the model already published indicating that miR159 impacts cell division and development via miR156/SPLs modules. In the present work, the Dual Luciferase Reporter Assay showed that SPL2 and SPL4 were silenced when agroinfiltrated with a miR156 precursor, meanwhile no interaction was detected with SPL13B, indicating that miR159 pathway controlling development and cell division involved only specific SPLs homologous genes. Most species also possess another miRNA, miR157, that has been described to be closely related to miR156 (Reinhart et al. 2002). miR157 has the same targets as miR156 and produces an over-expression phenotype similar to miR156 (Shikata et al. 2012). However, the function of miR157 is still unknown. We found that this miR157 was up regulated after genomic ‘shock’ in the presence of Phenanthrene. Hence, it is reasonable to speculate that this microRNA may also control plant development and cell division following genomic ‘shock’ in stressed conditions.

**Figure 9.**
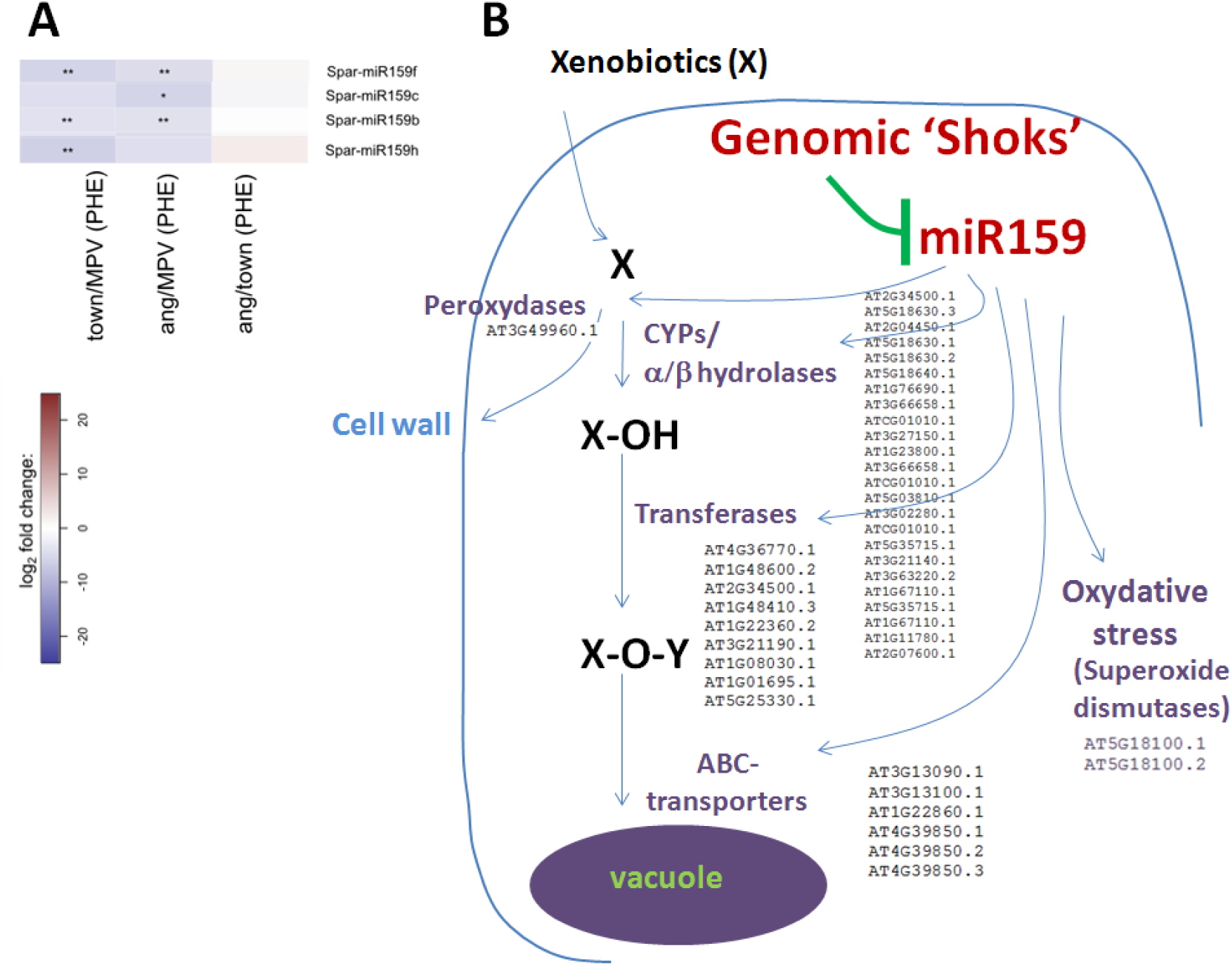
Negative impact of genomic ‘shock’ on *MIR*159 family and relaxation of the target xenome genes under xenobiotic induced stress. A: Heat map of almost DE miR159 genes depicting the ratio between the hybride (*S. x townsendii)* and the allopolyploide (*S. anglica)* and the MPV, respectively. B: the most genes with the highest prediction ranking scores (maximum expectation was set to 5.0 from psRNATarget; Dai *et al*. 2018), accession numbers were indicated in black) belong to the xenobiotic plant detoxification system and potentially relaxed upon MIR159 family genes inhibition. The xenobiotics are chemically modified using oxidation, reduction, or hydrolysis. Modified xenobiotics are conjugated to endogenous molecules. Transferases such glycosyltransferases (UGTs) transfer nucleotide-diphosphate-activated sugars such as UDP-glucose to low-molecular-weight substrates. The conjugated xenobiotic is transferred to the vacuole by ATP-binding cassette (ABC) transporters or to the cell wall.

The phenotype detected after screening FLAG_090F02 and FLAG_104E09 mutants suggested that plants lacking ath-MIR156e are sensitive to phenanthrene. The regulations between ath-miR156e and *SPL2* and *SPL4* were experimentally validated here, so down-regulation of ath-MIR156e (Spar-MIR156e homolog) is leading to up-regulations of *SPL* genes and phenanthrene sensitivity. Thus, we confirmed that phenanthrene induced ath-MIR156e (homologous to Spar-miR156e and j) acts on the down-regulation of negative stress tolerance regulators. Contrary to SALKseq_071642 harboring a T-DNA insertion in the predicted MIR156 like gene (homologous to Spar-miR156e and j), no inhibition upon treatment of high concentration of phenanthrene was observed. Indeed, many cases of the implication of lncRNAs in abiotic stress have been reported (Valadkhan & Valencia-Hipólito, 2015; Wang *et al*., 2017; Huanca-Mamani *et al*., 2018), through miRNA target gene mimicry, small RNAs precursor, or epigenetic modifications (histones or DNA methylation modifications). We did not validate any interaction between this putative miRNA and SPL genes, suggesting that this supposed novel miRNA precursor is not processed to mature miRNA in A*rabidopsis*.

Complementary functional validations are needed, to bring a broad picture of species-specific miRNAs involvement in *Spartina* xenobiotics tolerance. Such analyzes (for review see Eamens & Wang, 2011) in closely-related species are necessary to decipher miRNA functions in response to xenobiotics, especially for novel and specific *spartina* miRNAs. Using target mimicry (Franco-Zorrilla *et al*., 2007) or artificial miRNAs (Eamens *et al*., 2011) can be useful for such validations.

## Author contributions

A.E, A.S, A.C-R. and M.A. designed the experiments. A.E., O.L., R.M., L.T., L.S.P. and A.C.-R. performed the experiments. A.C-R., A.S., M.A. and A.E. analyzed data. A.C-R., M.A, A.S., and A.E. wrote the article.

## Acknowledgements

This work was supported by the Ministère de l’Enseignement Supérieur et de la Recherche, by the CNRS, and the Observatoire des Sciences et de l’Univers de Rennes (OSUR) and the International Associated Laboratory « Ecological Genomics of Polyploidy (LIA ECOGEN). This work was also partly granted by the University of Rennes 1 through the “Action inicitative Défis scientifiques 2018” awards to A.E., and the FASIC PROGRAM (Franco-Australian Hubert Curien Program) awards to A.E. The authors would like to acknowledge professor Peer Schrenk (university of Queensland) for valuable discussions, the Molecular Ecology Platform (UMR CNRS 6553 ECOBIO), the Human and Environmental Genomics Platform (Biogenouest) and the Genouest Bioinformatics Platform (Biogenouest).

## Supplementary data

**Supplemental Table S1.**
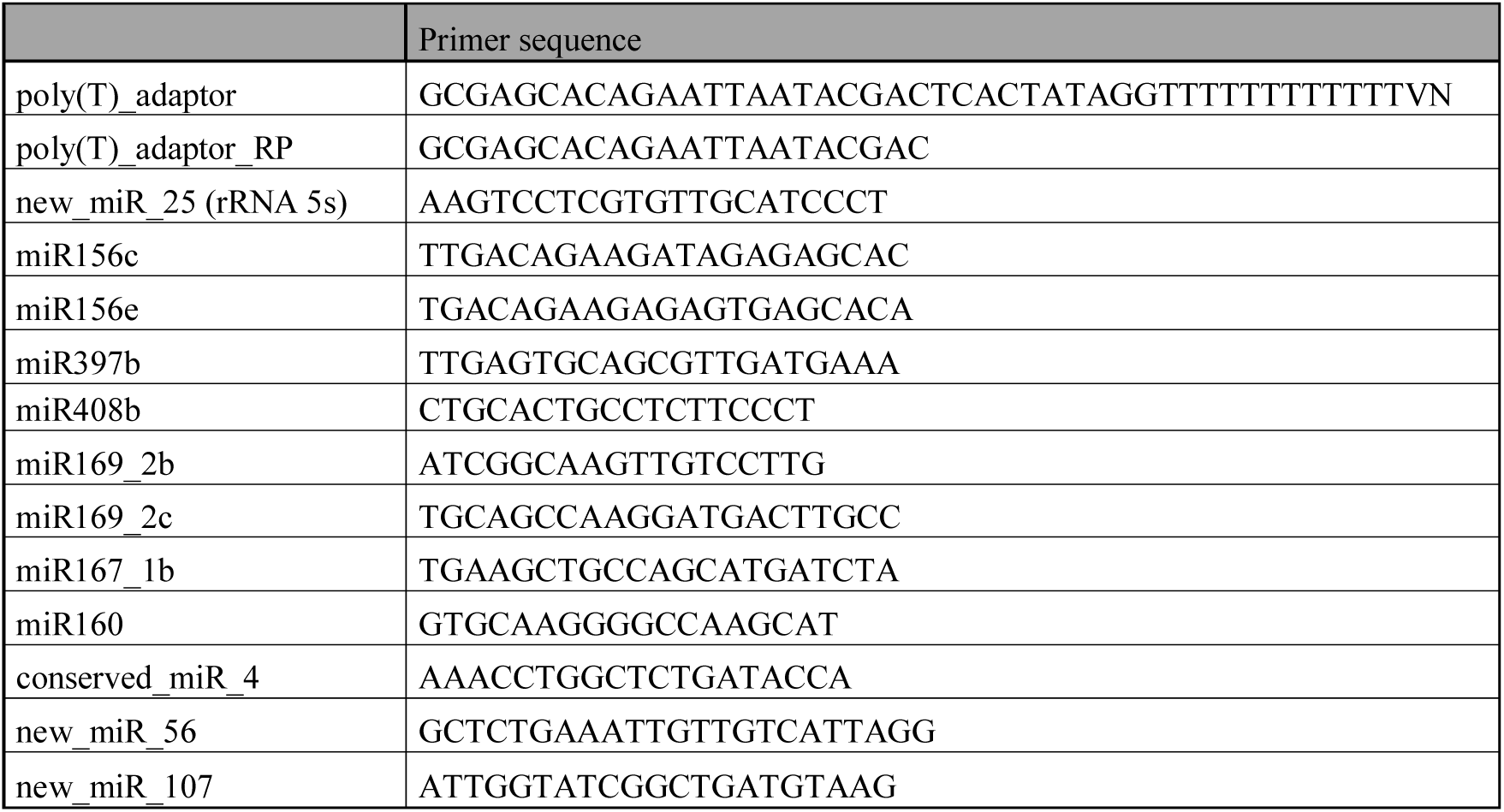
Primers used for RT-qPCR of miRNAs. The poly(T) adaptor used in reverse transcription of polyadenylated RNA is provided, as the universal poly(T) reverse primer sequence.

**Supplemental Table S2.** TAIR10 gene families potentially involved in xenobiotic detoxification (Edwards *et al*., 2011), or annotated as antioxidants enzymes and transcription factors (TFs) (https://www.arabidopsis.org/browse/genefamily/index.jsp)

**Supplemental Table S3.**
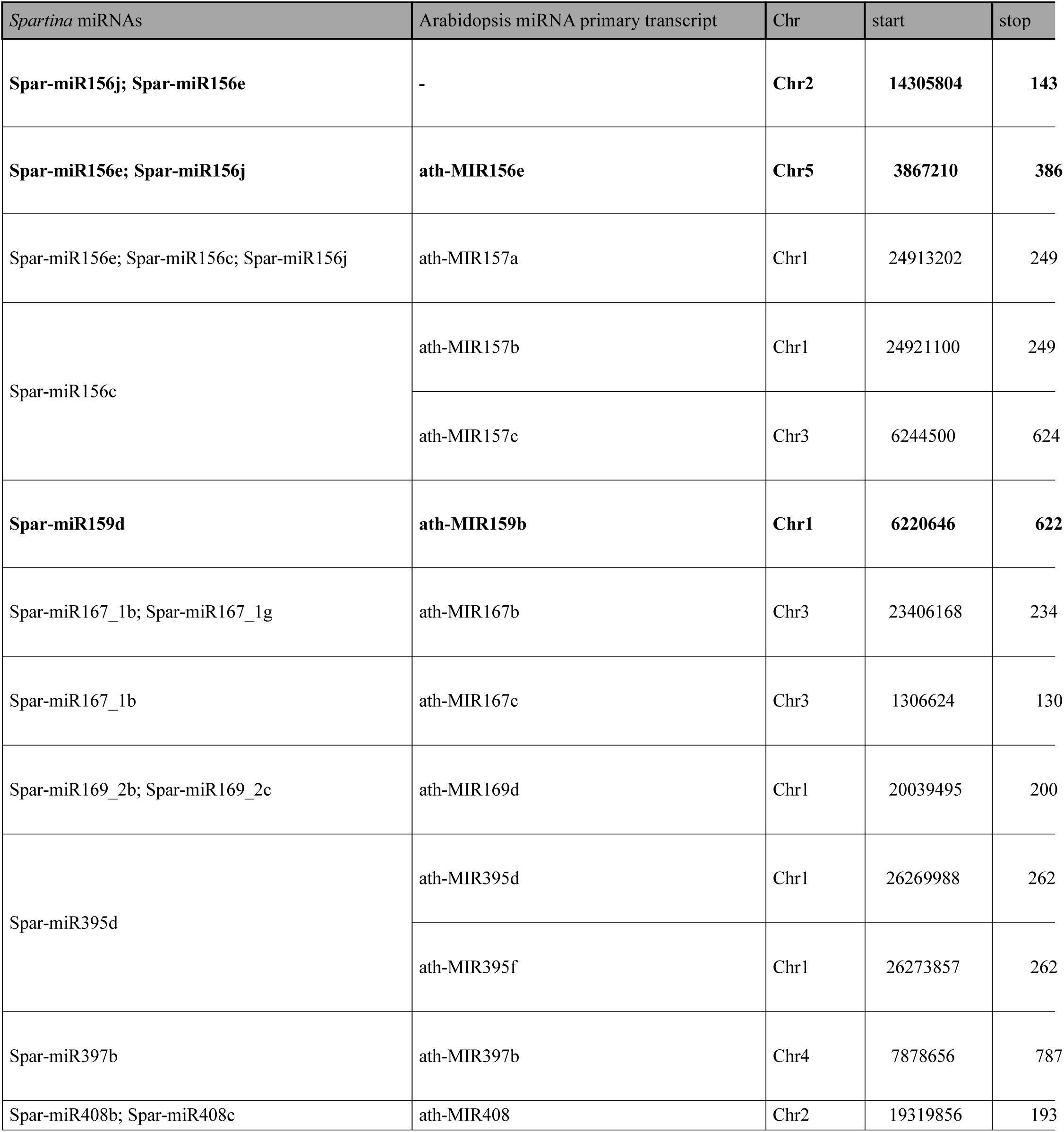
*Spartina* homologous *MIR* genes in *Arabidopsis* and corresponding T-DNA mutant lines. *Arabidopsis thaliana* T-DNA insertional mutants were obtained from the SALK, GABI-KAT and SAIL collections (Colombia (Col-0) genetic background), and from the FLAG collection (Wassilewskija (WS) background). The sequences used for mutants genotyping are indicated as LP and RP primers.

**Supplemental Table S4.**
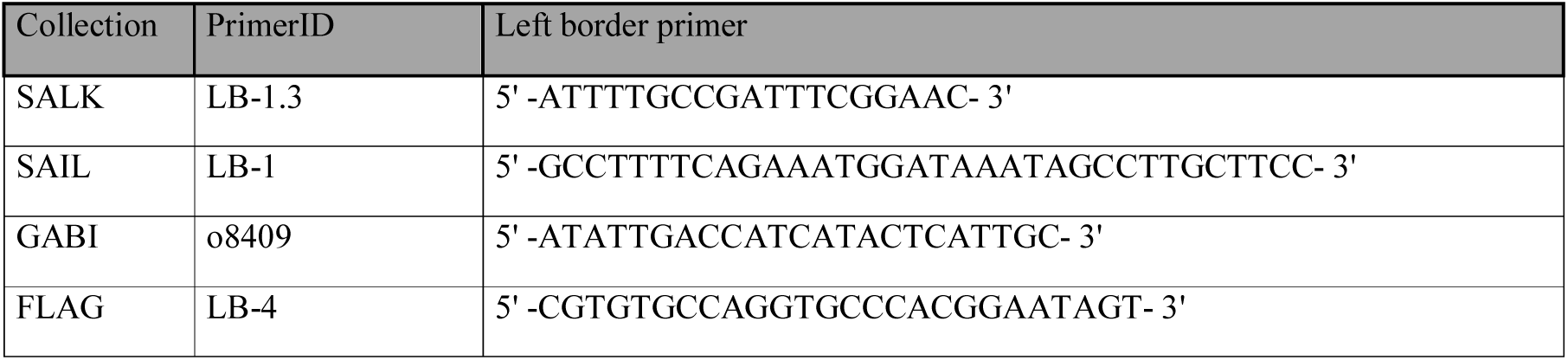
Left border primers used for T-DNA genotyping between mutant collections.

**Supplemental Table S5.**
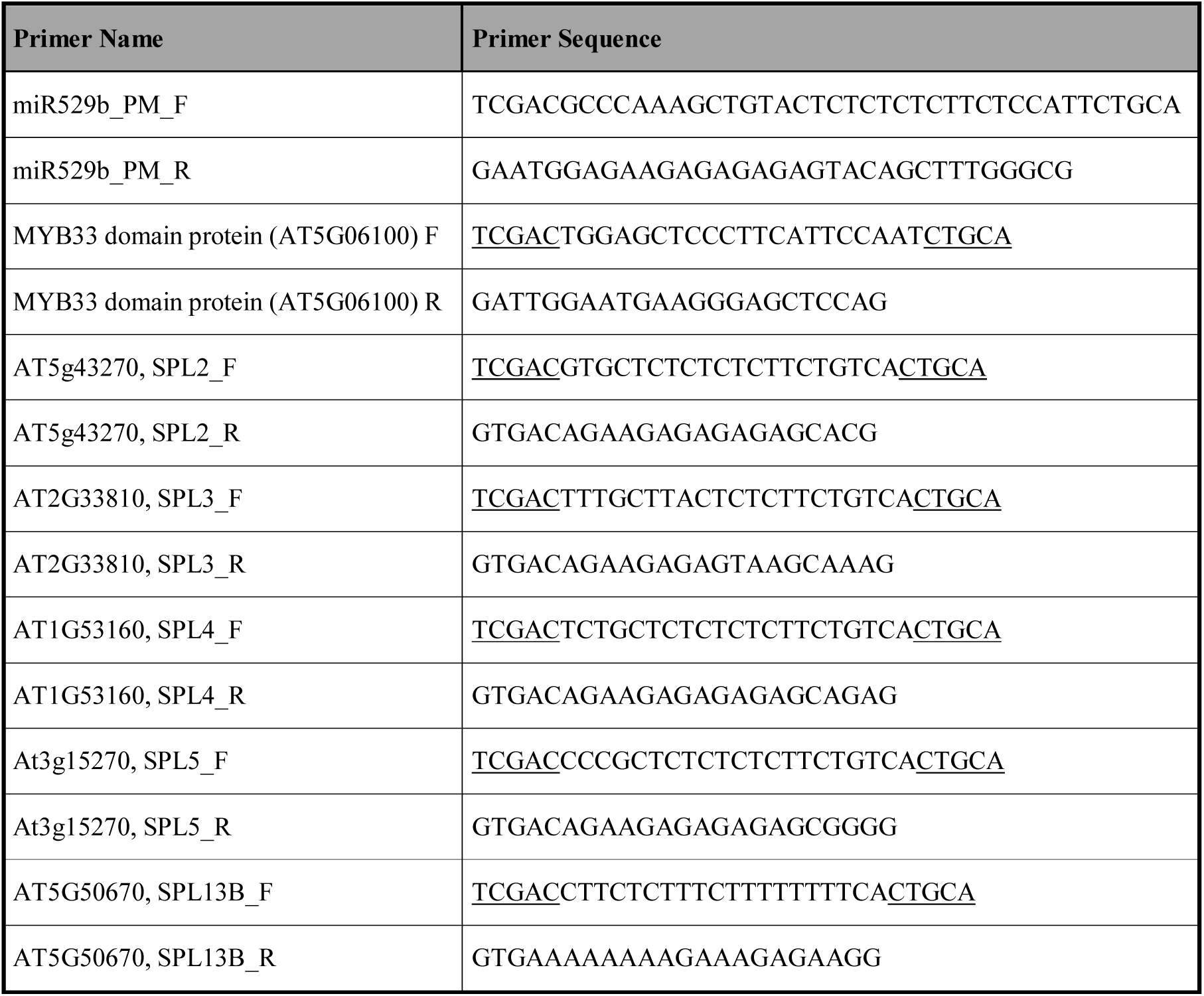
Target site adaptors used to amplify the target sequences used for the cloning in the plasmid pGrDL_SPb and in dual Luciferase assays.

**Supplemental Table S6.** Highly DE miRNA within *Spartina* species (log_2_ FC > 5 or < -5). About the total of 103 DE miRNAs identified 27 present major up or down-regulations in response to stress and represent candidates of interest in studying stress responsive miRNAs. Significant difference in expression are included: *: p.value < 0.05; **: p.value < 0.01.

| miRNA_name | *S. alterniflora* | sign | *S. maritima* | sign | *S. x townsendii* | sign | *S. anglica* | sign |
| --- | --- | --- | --- | --- | --- | --- | --- | --- |
| Spar-miR156c | -2,09 | NA | -6,07 | ** | 3,24 | ** | 3,19 | ** |
| Spar-miR156e | 5,04 | ** | 1,41 | NA | 1,08 | NA | -0,79 | NA |
| Spar-miR159h | 5,06 | ** | 3,15 | NA | -1,17 | NA | 1,07 | NA |
| Spar-miR159i | 7,54 | ** | 1,70 | NA | 0,37 | NA | 0,41 | NA |
| Spar-miR160 | 0,85 | NA | 1,34 | NA | 11,97 | ** | -3,50 | ** |
| Spar-miR395b | -6,91 | ** | -3,27 | NA | -2,08 | NA | 4,12 | * |
| Spar-miR395d | -1,01 | NA | -0,89 | NA | 8,54 | ** | 5,02 | ** |
| Spar-miR398 | 2,57 | NA | 5,11 | * | -4,36 | NA | 1,19 | NA |
| Spar-miR399 | 0,25 | NA | -0,84 | NA | 6,07 | ** | 2,16 | NA |
| Spar-miR408b | 3,70 | NA | 5,80 | ** | -2,20 | NA | 3,86 | NA |
| Spar-miR408c | 4,46 | NA | 5,91 | ** | -4,37 | NA | 3,34 | NA |
| Spar-miR529 | -5,03 | ** | -1,25 | NA | -1,63 | NA | 1,55 | NA |
| Spar-new_miR_110 | - | - | -0,20 | NA | 5,76 | ** | -0,32 | NA |
| Spar-new_miR_202 | -5,79 | ** | -0,38 | NA | -3,73 | * | 1,70 | NA |
| Spar-new_miR_271 | - | - | -0,42 | NA | -5,63 | ** | -3,20 | NA |
| Spar-new_miR_32 | - | - | -0,89 | NA | 8,03 | ** | 2,87 | NA |
| Spar-new_miR_360 | 1,65 | NA | -0,86 | NA | -3,80 | ** | 5,82 | ** |
| Spar-new_miR_362 | -2,00 | NA | 0,74 | NA | -5,47 | ** | 2,87 | NA |
| Spar-new_miR_384 | - | - | -0,30 | NA | 7,54 | ** | 8,99 | ** |
| Spar-new_miR_409 | - | - | - | - | 5,58 | ** | 0,78 | NA |
| Spar-new_miR_422 | -6,99 | ** | - | - | -7,34 | ** | -1,22 | NA |
| Spar-new_miR_434 | -1,05 | NA | -1,64 | NA | -5,09 | ** | 4,47 | ** |
| Spar-new_miR_468 | -7,48 | ** | - | - | -1,70 | NA | 2,57 | NA |
| Spar-new_miR_479 | 1,67 | NA | 0,39 | NA | 5,25 | * | -0,43 | NA |
| Spar-new_miR_5 | 5,28 | * | 1,13 | NA | 2,00 | NA | 1,30 | NA |
| Spar-new_miR_51 | - | - | -1,34 | NA | 5,24 | ** | -1,74 | NA |
| Spar-new_miR_69 | 0,39 | NA | - | - | 0,67 | NA | -6,77 | ** |

**Supplemental Table S7.**
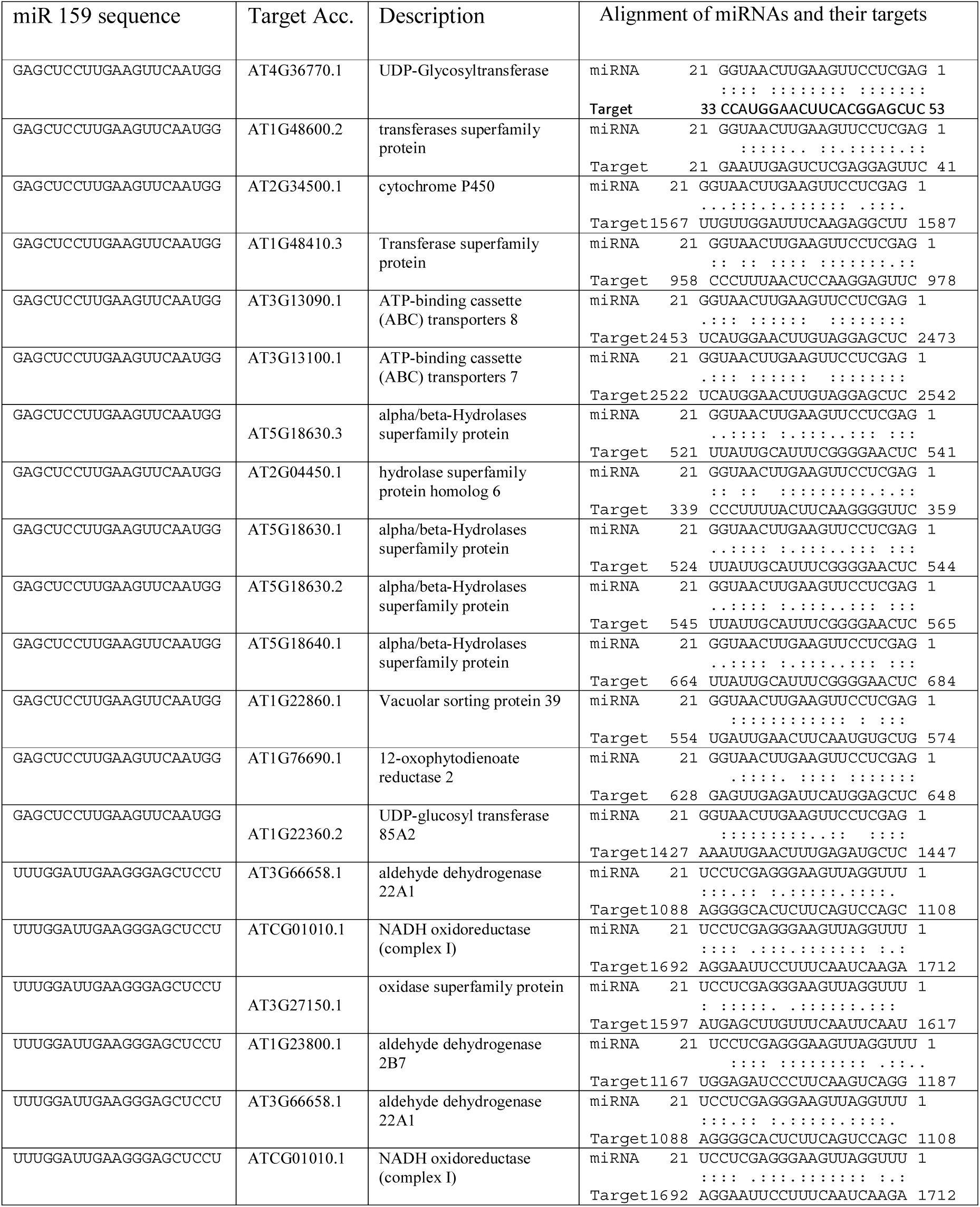
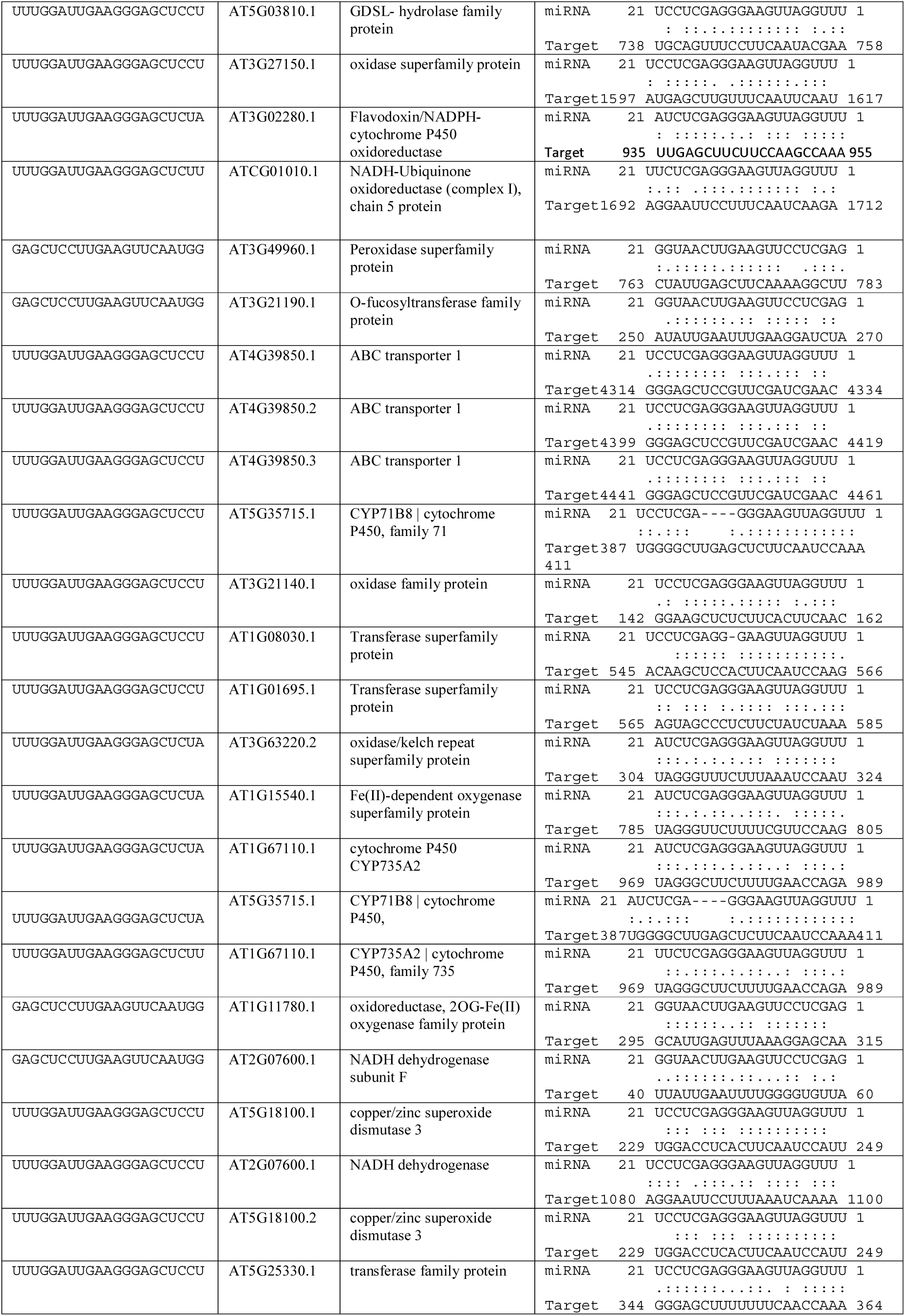
Target accessions with the highest prediction ranking scores (above 5.0 from psRNATarget; Dai et al. 2018) and alignment indicating high representativity of xenome products of almost all xenobiotics detoxification steps as shown in figure 9B. MIR159 family target also superoxide dismutases, known to be involved in ROS scavenging under xenobiotics induced stress.

**Supplemental Figure S1.**
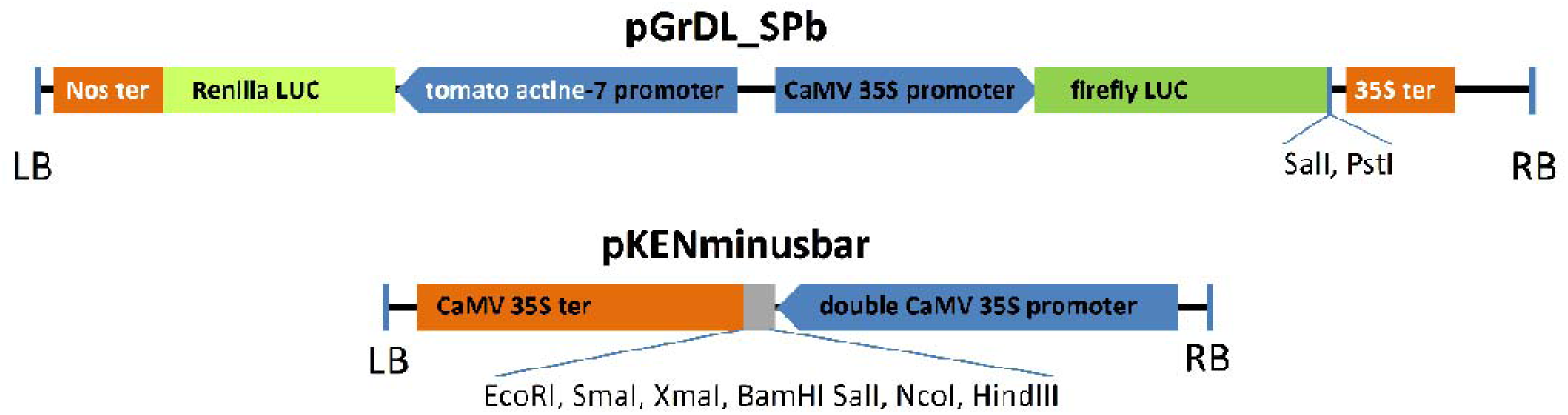
Plasmids used for Transient assay systems to validate computationally predicted targets of miRNA. The plasmid pGrDL_SPb contain the renilla LUC expression cassette which acts as an internal control to standardize expression between replicates, while the firefly LUC expression cassette containing the predicted target sequence cloned between SalI and PstI is used to report miRNA interaction. Expression plasmid pKENminusbar with multiple cloning sites is used for miRNA precursor insertion.

**Supplemental Figure S2.**
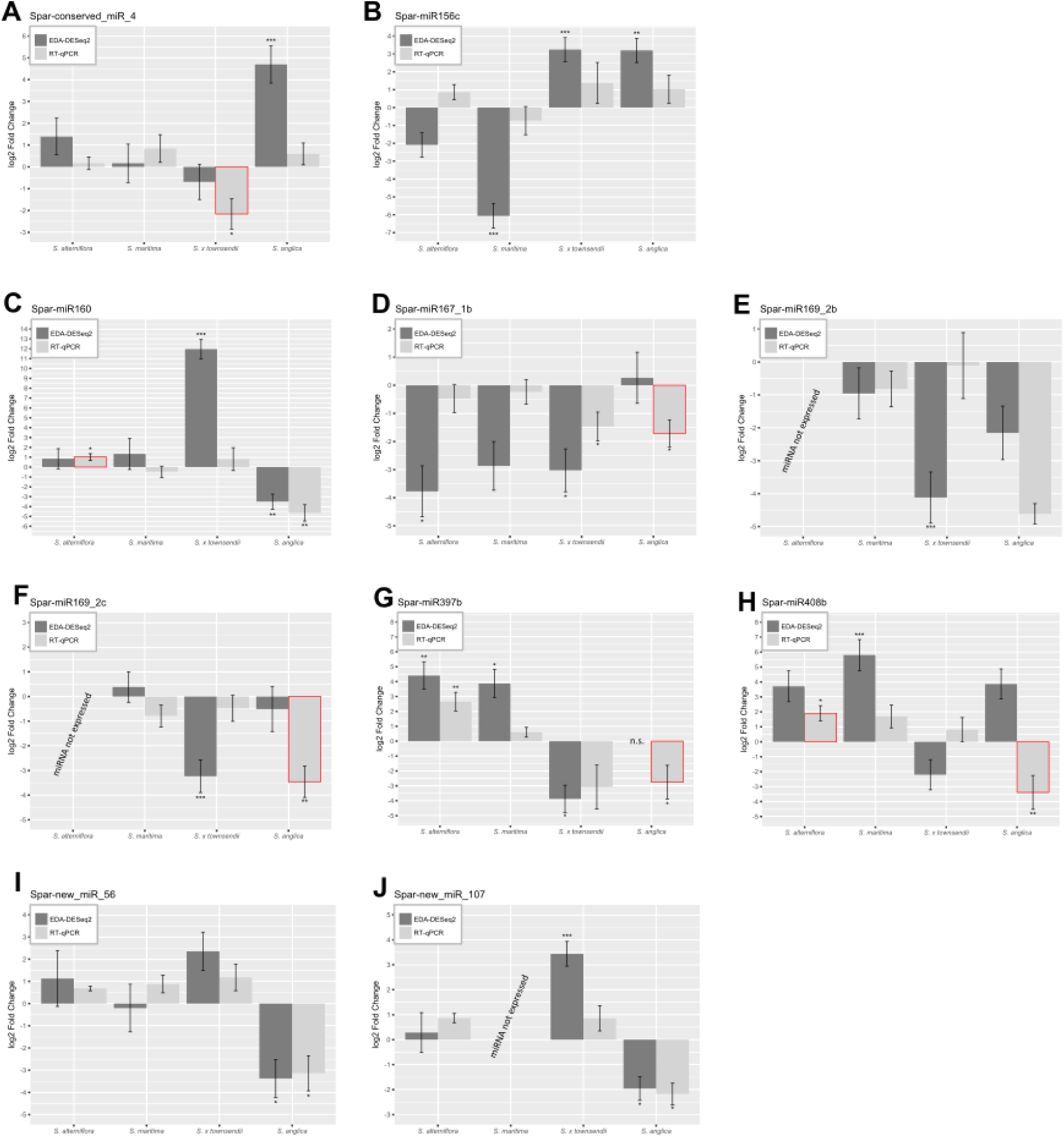
Micro-RNA expression log_2_ Fold Changes obtained by RT-qPCR for 11 RNA-Seq DE expressed miRNAs. Results are confronted among *Spartina* species with EDA-Seq and DESeq2 *in silico* data (*: p.value < 0.5; **: p.value < 0.1; **: p.value < 0.01). Here, 10 significant regulation profiles (bars surrounded in red Figure 3) were newly identified by RT-qPCR.

**Supplemental Figure S3.**
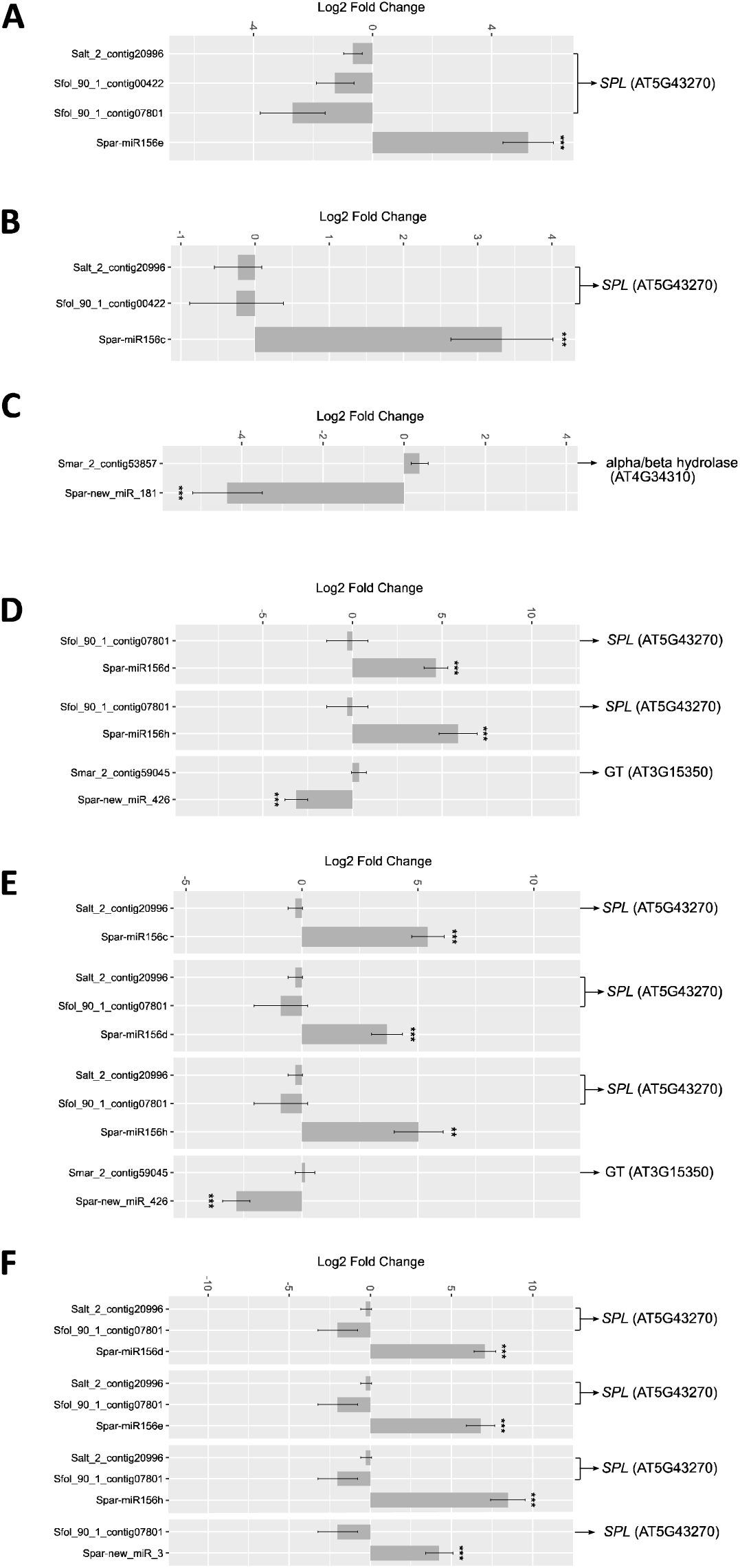
Differentially expressed miRNAs, based on intraspecific comparisons in response to phenanthrene in *S. alterniflora* (A), *S. x townsendii* (B), *S. anglica* (C) and interspecific comparisons between PAH tolerant species *S. anglica* (D), *S. x townsendii* (E) and *S. alterniflora* (F) compared to the PAH sensitive *S. maritima*. Opposite expression patterns of miRNA target genes annotated among phenanthrene-responsive candidates (*SPL*, α/β hydrolases and *GT*) in *Arabidopsis* are provided.

